# Multi-Substrate Specificity of Isoflavone hydroxylases (GmIFH) Drive Isoflavonoid Diversification in Soybean

**DOI:** 10.64898/2026.05.05.722824

**Authors:** Praveen Khatri, Tim McDowell, Frédéric Marsolais, Justin Renaud, Sangeeta Dhaubhadel

## Abstract

Isoflavone hydroxylases (IFHs, CYP81E) convert isoflavone aglycones into their respective hydroxylated intermediates, which direct legume isoflavones into specialized defense pathways. In soybean, their functions have been studied mostly in the context of the daidzein-derived glyceollin biosynthesis. Here we combine metabolomics-guided feature mining, phylogenetic analysis, heterologous enzymology, structural elucidation, and *in planta* metabolite validation to determine the functional landscape of the soybean IFH family. Analysis of a soybean isoflavonoid-enriched metabolomic dataset revealed unidentified hydroxyisoflavone features that co-accumulated with glyceollins, indicating branch chemistry that is not well-recognized. The systematic characterization of the repertoire of soybean CYP81E has demonstrated that 9 out of 11 GmIFHs are catalytically active and collectively span both 2′- and 3′- hydroxylation of the major soybean isoflavone aglycones. Among them, GmIFH9A showed broad substrate scope and regioselectivity, yielding canonical and previously unknown hydroxylated isoflavone products. NMR and LC-MS/MS were used to identify and validate the hydroxylated isoflavone products as 2′-hydroxyglycitein and 2′-hydroxyformononetin, whose presence was also confirmed in soybean roots, thus confirming two of the hidden soybean isoflavonoid network metabolites. Kinetic studies also indicated that, although the majority of GmIFHs prefer daidzein and genistein as substrates, a few isoforms are active towards methoxylated isoflavones as well, indicating functional divergence in this expanded family. Our findings collectively redefine soybean IFHs as a multi-functional enzyme module that expands the hydroxyisoflavone chemical space and reveals new biosynthetic entry points beyond canonical glyceollin pathway.

## Introduction

Legume plants predominantly accumulate five isoflavone aglycones namely: daidzein, genistein, glycitein, formononetin, and biochanin A (Anguraj Vadivel et al., 2019; Liu et al., 2003) whose differential hydroxylation by isoflavone hydroxylases (IFHs) on the B-ring generates a structurally diverse repertoire of hydroxyisoflavones. These hydroxylated isoflavones are among the earliest metabolites to accumulate following biotic or abiotic stress and undergo further enzymatic conversion into multiple classes of phytoalexins, including glyceollin, vestitol, medicarpin, maackiain, pisatin, coumestans, kievitone, and rotenoids (Clemens et al., 1993; Liu *et al*., 2003; Sukumaran et al., 2018).

IFHs belong to the CYP81E subfamily of cytochrome P450 monooxygenases (Akashi et al., 1998) and are of two major types based on their regioselectivity: isoflavone 2′-hydroxylases (I2′Hs) and isoflavone 3′-hydroxylases (I3′Hs) (Figure 1). I2′Hs have been characterized in many legumes including *Glycyrrhiza echinata* (Akashi *et al*., 1998), *Lotus japonicus* (Shimada et al., 2000), *Medicago truncatula* (Liu et al., 2003), and *Pueraria mirifica* (Suntichaikamolkul et al., 2022), where they are reported to hydroxylate major isoflavones (except glycitein) to generate precursors for phytoalexins such as vestitol, medicarpin, and miroestrol. I3′Hs convert biochanin A and formononetin into 3′-hydroxylated intermediates that feed into maackiain and pisatin biosynthesis in chickpea (*Cicer arietinum*) and pea (*Pisum sativum*) (Banks and Dewick, 1982; Clemens *et al*., 1993; Overkamp et al., 2000)

**Figure 1.**
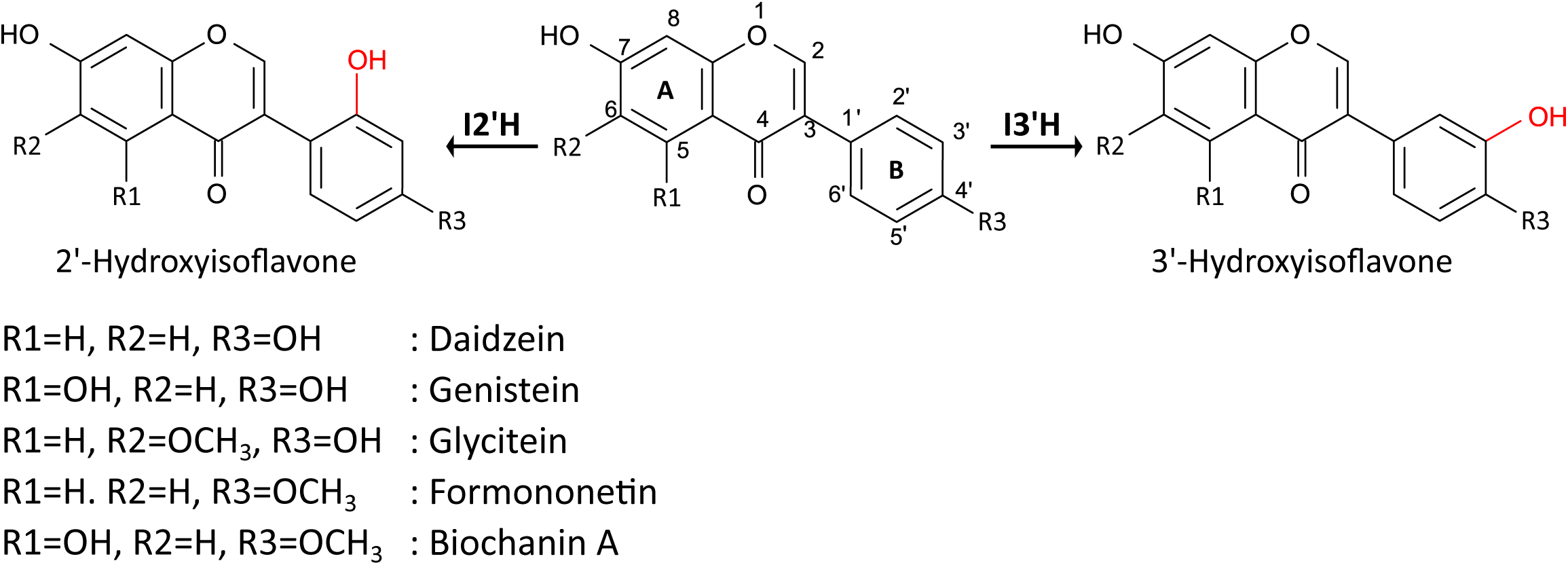
Chemical structure and numbering of isoflavone aglycones. Different isoflavones aglycons are shown by type of substitution in positions R1-R3. The reactions catalysed by isoflavone 2′-hydroxylase (I2′H) and isoflavone 3′-hydroxylase (I3′H) are shown, with the respective hydroxylation sites on the B ring indicated in red.

In soybean, IFHs catalyze the conversion of daidzein and genistein into their respective 2′-hydroxylated derivatives (Fischer et al., 1990; Simons et al., 2011). Among these, 2′-hydroxydaidzein serves as the key precursor for glyceollin biosynthesis, establishing IFH activity as indispensable for the formation of the primary soybean phytoalexin. Three CYP81E family members CYP81E11, CYP81E12, and CYP81E18 which encode I2′Hs that catalyse the enzymatic reactions to convert daidzein and genistein to their respective hydroxyisoflavones have been reported thus far (Uchida et al., 2015). Despite the evidence from other legumes that IFHs can generate structurally diverse hydroxylated intermediates via regioselective hydroxylation, functional studies of soybean IFHs have remained largely restricted to daidzein-centered glyceollin biosynthesis. Additionally, although untargeted metabolomics has consistently revealed features indicative of hydroxylated isoflavones in soybean tissues, the structures of many of these metabolites remain unresolved, and the specific enzymes responsible for their biosynthesis in soybean have yet to be identified.

In this study, we undertook a metabolomics-guided enzymology framework to (i) systematically characterize the entire soybean CYP81E repertoire, (ii) determine substrate scope and kinetic preferences of GmIFHs, and (iii) use enzymatically produced standards to identify unknown metabolite features in soybean roots. We confirm for the first time the presence of 2′-hydroxyglycitein and 2′-hydroxyformononetin in soybean roots, revealing possibility of presence of unrecognized branches of soybean isoflavonoids parallel to the well-established daidzein-centred glyceollin pathway. Collectively, our results reveal previously unrecognized metabolic pathways derived from glycitein and formononetin, the diversification and functional roles of the soybean IFH family, and new biosynthetic entry points for engineering isoflavonoids to improve plant resilience and human health.

## Results

### Mining of isoflavone-related metabolite features in soybean

To uncover previously unreported isoflavone-derived metabolites in soybean, we revisited our LC-MS dataset generated from the *GmMYB176-GmbZIP5* overexpressing hairy-roots, where coordinated expression of these transcriptional regulators increased isoflavonoid accumulation (Anguraj Vadivel *et al*., 2021). Targeted feature mining using predicted adduct masses (Supplementary Table 1) for the five major isoflavone aglycones (daidzein, genistein, glycitein, formononetin, and biochanin A) and their hydroxylated derivatives identified multiple significantly induced LC-MS features (log2 fold change > 1, FDR < 0.05) that remained unassigned (Figure 2A, Supplementary Table 1). Several of these induced ions were consistent with putative hydroxyformononetin- and hydroxyglycitein-like masses detected in both positive and negative ionization modes, suggesting the presence of novel hydroxylated isoflavone intermediates in soybean.

**Figure 2.**
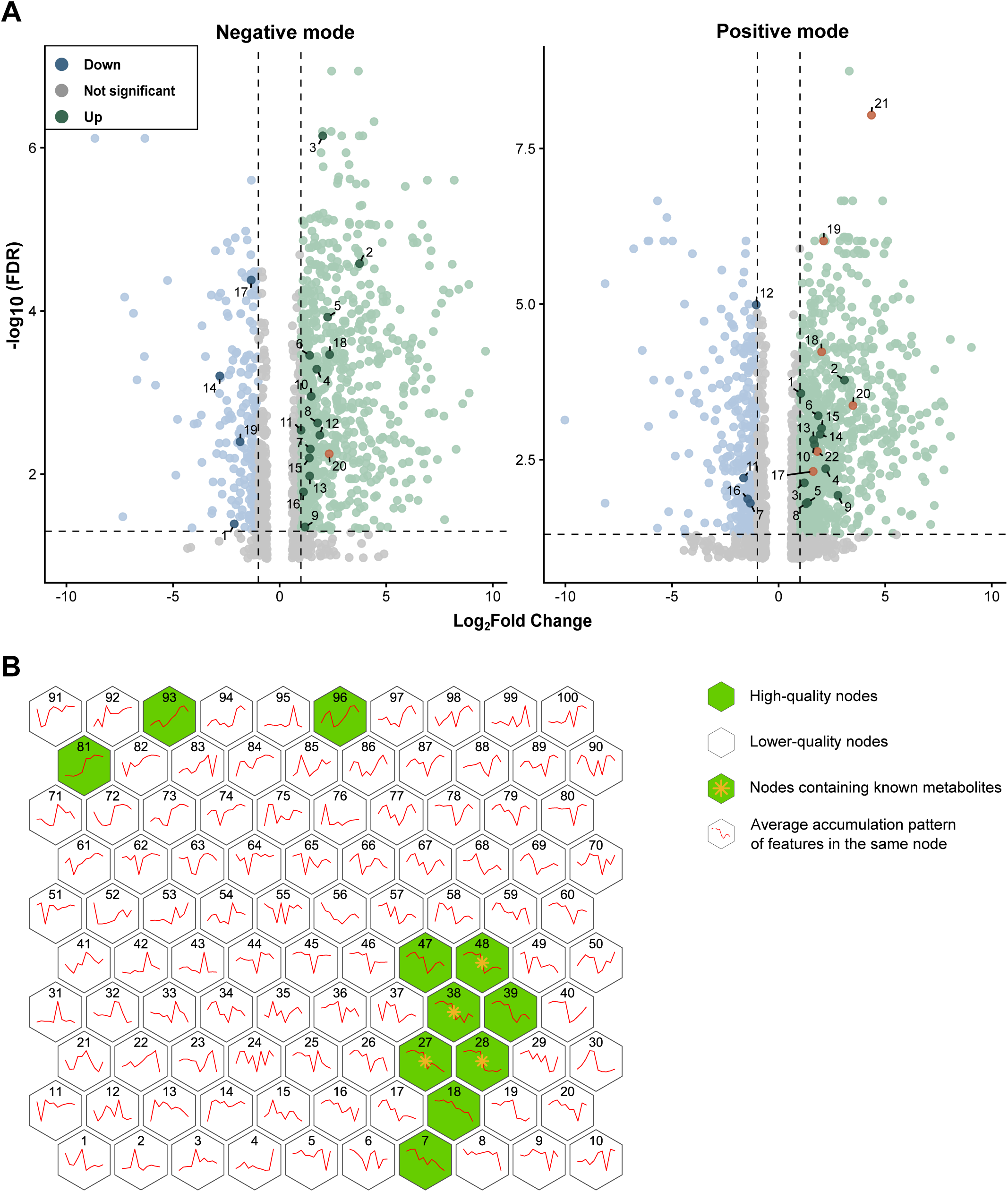
Analysis of 2’hydroxyisoflavones in soybean hairy roots overexpressing GmMYB176 and GmbZIP5. **(A)** Volcano plot showing the differentially accumulated metabolites in negative and positive modes. Highlighted dots show differentially accumulated hydroxyisoflavone features. Mustard dots show previously identified isoflavone features. **(B)** Self organizing map (10x10) showing nodes with number of metabolites co-accumulating with most similar accumulation profile. The reported isoflavone metabolites from Anguraj Vadivel et al., 2021 are represented by yellow stars (⁎). Green nodes signify the highest quality nodes, white nodes are of lower quality.

To further link these candidate features to the soybean phytoalexin response, we constructed a self-organizing map (SOM) to identify metabolites that co-accumulate with glyceollin (*m/z* 339.12, RT 3.5 min). This analysis positioned glyceollin within the node 48 which also contained a hydroxydaidzein-like feature consistent with its established association with the glyceollin pathway, along with hydroxyformononetin and hydroxyglycitein like features indicating coordinated accumulation within the same metabolome. More broadly, the isoflavonoid-related characteristics were localized in a few highlighted nodes of the SOM as opposed to scattered throughout the map, with a few high-quality nodes clustered together around node 48 (Figure 2B, Supplementary Figure 1). This local clustering correlates with common patterns of abundance of these metabolites and confirms the existence of a coordinated isoflavonoid response unit that is linked to the accumulation of glyceollin. Nodes clustered with 48 (such as 27, 28 and 38) had isoflavone-metabolites that were reported in the metabolomics dataset earlier (Anguraj Vadivel *et al,* 2021), suggesting that the induced soybean metabolome is structured into a few, related, yet distinct, isoflavonoid accumulation patterns. Together, these observations motivated us for a metabolomics-guided enzymology strategy to identify the enzyme(s) responsible for generating these hydroxylated isoflavones and to produce enzymatic reference standards for definitive structural assignment. Given that CYP81E enzymes in legumes commonly function as isoflavone 2′- and/or 3′-hydroxylases, we next focused on systematic characterization of the soybean CYP81E repertoire as candidate enzymes responsible for generating these hydroxylated compounds.

### Identification, phylogeny and sequence analyses of CYP81 candidates in soybean

To define candidate enzymes, we constructed a maximum-likelihood phylogenetic tree using 40 functionally characterized CYP81 proteins from 22 diverse plant species (Supplementary Table 2), together with the 12 annotated soybean CYP81 members (Khatri et al 2022). The analysis revealed that, with the exception of Glyma.11G093100.1, which clustered with CYP81 enzymes involved in the terpenoid (Wang et al., 2021), xanthone (El-Awaad et al., 2016), and quinone biosynthesis pathway (Ren et al., 2023), all soybean CYP81 members grouped within the CYP81s clade previously reported for their IFH activity. We therefore designated the remaining eleven soybean CYP81 members as the putative GmIFHs and named them based on their respective chromosomal locations (Table 1). Within the phylogeny, GmIFH8A, GmIFH11, and GmIFH16 grouped with I3′H from *M. truncatula* (CYP81E9) whereas the remaining eight GmIFHs clustered with I2′H from *L. japonica* (LjCYP-2), *G. echinata* (CYP81E1), *M. truncatula* (CYP81E7) and *P. mirifica* (CYP81E63) (Figure 3). All GmIFHs encode proteins of an average length of approximately 500 amino acids with a predicted molecular weight of 57.31 to 65.04 kDa. The predicted isoelectric points (pIs) ranged from 8.06 to 9.29, indicating an overall basic nature across all GmIFH proteins and were predicted to localize to the endoplasmic reticulum (Table 1). Multiple sequence alignment further revealed that all GmIFHs retain the canonical cytochrome P450 structural features, including the N-terminal proline-rich region, the I-helix oxygen-binding motif, the K-helix ETLR and PERF motifs, and the C-terminal heme-binding loop (Supplementary Figure 2). While sequence identity varied across the different enzyme types, particularly between I2′H and I3′H like members (Supplementary Figure 3), conservation of these hallmark motifs supported the enzymatic competence of all candidates.

**Figure 3.**
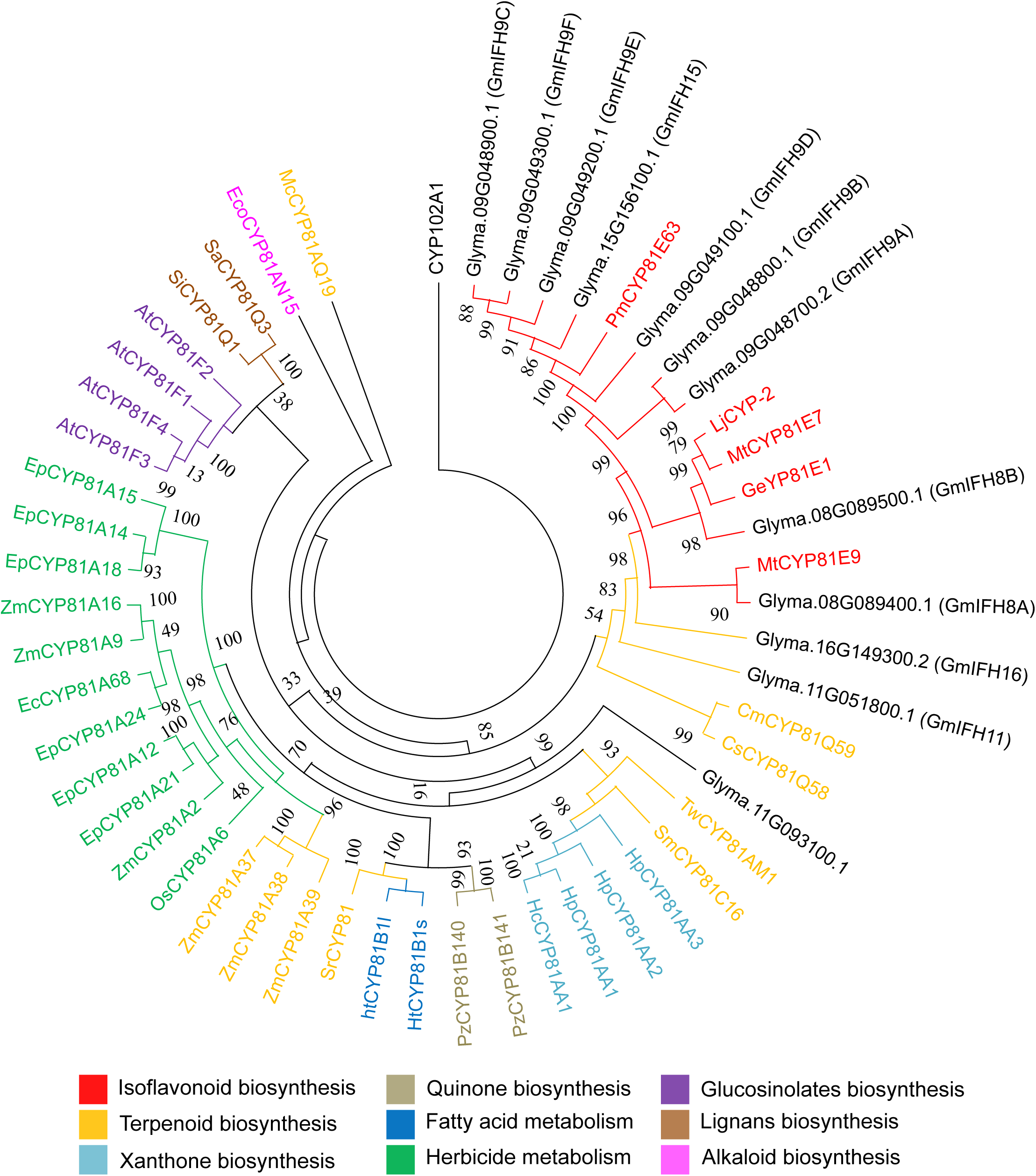
Phylogenetic analysis of CYP81 proteins. Amino acid sequences of characterized CYP81s from 22 plant species and soybean CYP81s were aligned in ClustalO and a phylogenetic tree was constructed using a maximum likelihood method in IQ-Tree with best fit model with a bootstrap value set to 1,000. CYP102A1 indicates an out-group used to root the tree. Different colors indicate the functional classes to which CYP81s belong. At, *Arabidopsis thaliana;* Mt, *Medicago truncatula;* Ge, *Glycyrrhiza echinate;* Ht, *Helianthus tuberosus;* Lj, *Lotus japonicus;* Pm, *Pueraria mirifica;* Cm*, Cucumis melo;* Cs: *Cucumis sativus;* Tw, *Tripterygium wilfordii;* Sm, *Salvia miltiorrhiza;* Hp, *Hypericum perforatum;* Hc, *Hypericum calycinum,* Pz, *Plumbago zeylanica;* Sr, *Stevia rebaudiana;* Zm, *Zea mays,* Ec, *Echinochloa phyllopogon ;* Os, *Oryze sativa,* Ec, *Echinochloa crus-galli;* Si, *Sesamum indicum;* Sa, *Sesamum alatum;* Eco*, Erythroxylum coca;* Mc*, Momordica charantia*

**Table 1.** Detailed information of *GmIFH*s and their physiochemical properties.

| Name | Locus ID | Genomic location | Molecular weight (kDa) | Predicted isoelectric point (pI) | CDS length (bp) | Splice variants | Number of introns |
| --- | --- | --- | --- | --- | --- | --- | --- |
| GmIFH8A | *Glyma.08G089400* | Chr08 : 6764477..6767073 | 57.31 | 8.53 | 1503 | 1 | 2 |
| GmIFH8B | *Glyma.08G089500* | Chr08 : 6770389..6772737 | 57.49 | 8.99 | 1509 | 1 | 2 |
| GmIFH9A | *Glyma.09G048700* | Chr09 : 4180465..4183894 | 58.20 | 8.06 | 1506 | 1 | 2 |
| GmIFH9B | *Glyma.09G048800* | Chr09 : 4184294..4186825 | 58.01 | 9 | 1497 | 1 | 2 |
| GmIFH9C | *Glyma.09G048900* | Chr09 : 4193594..4195649 | 57.55 | 8.7 | 1506 | 2 | 1 |
| GmIFH9D | *Glyma.09G049100* | Chr09 : 4234839..4236838 | 57.91 | 9.03 | 1512 | 1 | 1 |
| GmIFH9E | *Glyma.09G049200* | Chr09 : 4253575..4256263 | 57.35 | 8.81 | 1497 | 1 | 1 |
| GmIFH9F | *Glyma.09G049300* | Chr09 : 4259304..4261332 | 57.43 | 8.82 | 1503 | 1 | 1 |
| GmIFH11 | *Glyma.11G051800* | Chr11 : 3886032..3888141 | 56.23 | 9.26 | 1491 | 1 | 1 |
| GmIFH15 | *Glyma.15G156100* | Chr15 : 13074247..13076579 | 57.65 | 8.57 | 1509 | 1 | 1 |
| GmIFH16 | *Glyma.16G149300* | Chr16 : 31186383..31190129 | 65.04 | 9.29 | 1692 | 1 | 2 |

### Functional characterization of *GmIFHs* to produce hydroxyisoflavones

To test whether candidate GmIFHs could generate the unassigned metabolites detected by metabolomics, we heterologously expressed GmIFHs in yeast and performed microsomal enzyme assays using individual isoflavone aglycones as substrates. Of the eleven GmIFHs tested, nine produced detectable hydroxylated products, whereas GmIFH11 and GmIFH16 showed no activity with any substrate under the conditions examined. When glycitein was used as substrate, six GmIFHs (GmIFH9A, GmIFH9C, GmIFH9D, GmIFH9E, GmIFH9F and GmIFH15) generated a product detected by HPLC-UV at retention time 11.33 min. Independent LC-MS analysis of the same reactions revealed a corresponding ion at *m/z* 301.07 ([M+H]⁺), matching the hydroxyglycitein-like feature previously observed in the untargeted metabolomics analysis (Figure 4; Supplementary Figure 4). Purification of this enzymatic product followed by NMR confirmed this product as 2′-hydroxyglycitein (Table 2, Figure 4D, Supplementary Figure 5-7), thereby resolving the chemical identity of a previously unassigned soybean metabolite. In parallel, enzymatically produced 2′-hydroxydaidzein and 2′-hydroxygenistein were also purified and analyzed by NMR to support comparative spectral assignment of 2′-hydroxyglycitein (Supplementary Figure 8-13). Similarly, enzymatic assays using formononetin yielded a hydroxylated product detected by HPLC–UV (RT 4.24 min) with a corresponding LC-MS signal at *m/z* 285.076 ([M+H]⁺), consistent with a hydroxyformononetin derivative observed in metabolomics analysis. Co-elution and mass spectral matching with the reaction product of the previously characterized *M. truncatula* I2′H (CYP81E7) confirmed this compound as 2′-hydroxyformononetin (Figure 4, Supplementary Figure 4).

**Figure 4.**
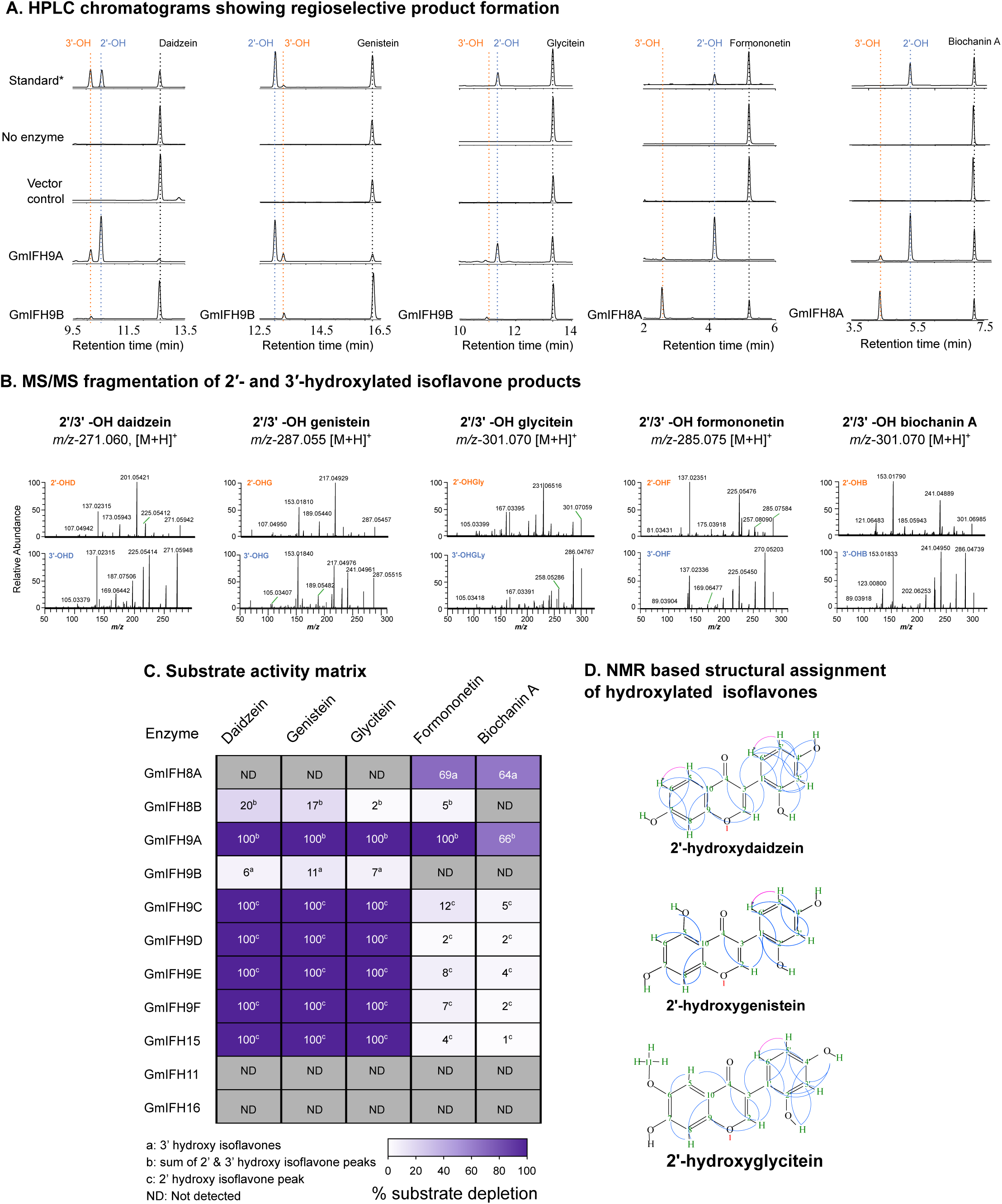
Functional characterization of soybean GmIFHs reveal broad substrate scope and distinct 2′- and 3′-hydroxylase activities toward isoflavones. **(A)** Representative HPLC chromatograms showing product formation from each substrate. GmIFH9A generated both 2′- and 3′-hydroxylated products from all five substrates. In contrast, GmIFH9B produced only 3′-hydroxylated products from daidzein, genistein, a putative 3′-hydroxylated product from glycitein, whereas GmIFH8A produced 3′-hydroxylated products from formononetin and biochanin A. Chromatograms include the indicated controls and representative enzyme reactions, with peak assignments based on retention time comparison with standards and LC-MS/MS analysis. **(B)** LC-MS/MS fragmentation analysis of hydroxylated products. Product identities were assigned based on diagnostic fragmentation patterns together with chromatographic behavior; the assignment of 3′-hydroxyglycitein is putative and based on accurate mass, LC-MS/MS fragmentation, and elution behavior relative to the confirmed 2′-hydroxyglycitein isomer. **(C)** Enzyme activity matrix summarizing the relative conversion of the five substrates by the indicated GmIFH isoforms. Activity was estimated from substrate depletion, and values represent the percentage of substrate converted under the assay conditions. **(D)** NMR-based structural assignment of hydroxylated isoflavones. HMBC (blue) and COSY (pink) correlations supporting the structural assignments of 2′-hydroxydaidzein, 2′-hydroxygenistein, and 2′-hydroxyglycitein are shown. These data were used to confirm hydroxylation position in compounds for which authentic standards were not available. Only diagnostic correlations relevant to structure validation are presented; full 1D and 2D NMR datasets are provided in the Supplementary Data. Standard* indicates an authentic standard commercially available, or an enzymatically produced compound that was structurally verified by NMR. Dotted lines indicate substrate and product peaks. 2’-OHD: 2’-hydroxydaidzein, 3’-OHD: 3’-hydroxydaidzein, 2’-OHG: 2’-hydroxygenistein, 3’-OHG: 3’-hydroxygenistein, 2’- OHGly: 2’-hydroxyglycitein, 3’-OHGly: 3’-hydroxyglycitein, 2’-OHF: 2’-hydroxyformononetin, 3’-OHF: 3’-hydroxyformononetin, 2’-OHB: 2’-hydroxybiochanin A, 3’-OHB: 3’-hydroxybiochanin A.

**Table 2.**
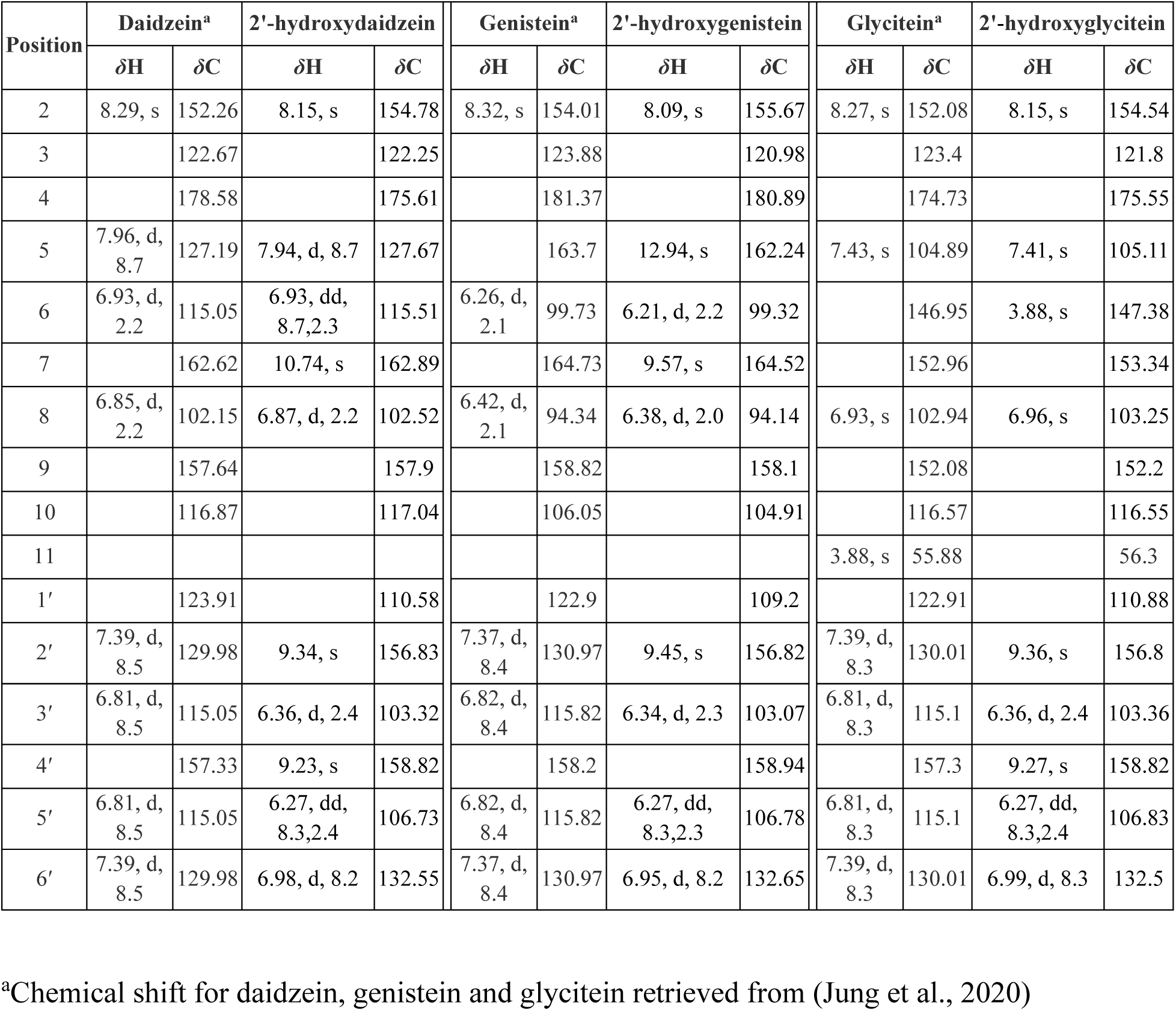
^1^H-NMR (*δ*_H_ in ppm, *J* in Hz) and ^13^C-NMR (*δ*_C_ in ppm) chemical shifts of daidzein, glycitein, and genistein and their 2’-hydroxy forms.

Among the enzymes tested, GmIFH9A exhibited the broadest catalytic scope, catalyzing both 2′- and 3′-hydroxylation reactions across all five major soybean isoflavone aglycones, including daidzein, genistein, glycitein, formononetin, and biochanin A, however, the assignment of 3′-hydroxyglycitein remains putative, based on accurate mass, LC-MS/MS fragmentation, and chromatographic behavior. (Figure 4c, Supplementary Figure 14-16). This dual-regioselective and multi-substrate activity directly accounts for the formation of both 2′-hydroxyglycitein and 2′-hydroxyformononetin and distinguishes GmIFH9A from other members of the GmIFH family.

In addition to resolving hydroxyglycitein and hydroxyformononetin, enzyme assays revealed broader hydroxylation activity across the GmIFH family. GmIFH9C, GmIFH9D, GmIFH9E, GmIFH9F and GmIFH15 primarily catalyzed 2’ hydroxylation of the isoflavones daidzein, genistein and glycitein and to a lesser extent of formononetin and biochanin A; consistent with canonical I2′H activity associated with glyceollin biosynthesis pathway. GmIFH9B functioned as an I3’H and converted daidzein, genistein and glycitein to their respective 3’-hydroxylated forms while the glycitein-derived 3′-hydroxylated product assignment remains putative based on LC-MS/MS and chromatographic behavior. GmIFH8A showed 3’-hydroxylase activity towards formononetin and biochanin A, thus showing I3’H type specificity. Whereas, GmIFH8B displayed weak catalytic activity, producing low levels of both 2′- and 3′-hydroxylated products from daidzein and genistein, while showing limited detectable activity toward glycitein and formononetin under the conditions tested (Figure 4, Supplementary Figure 4).

### Kinetic properties of GmI2′Hs

To evaluate whether the enzymatic formation of 2′-hydroxyglycitein and 2′-hydroxyformononetin is kinetically supported, we performed steady-state kinetic analyses on six GmIFHs exhibiting robust 2′-hydroxylation activity *in vitro*. Across the enzyme set, daidzein showed the most consistently favorable binding with *K*_m_ values in the low micromolar range (2.61-7.61 µM) for all six enzymes reflecting the established role of GmIFHs in supplying precursors for glyceollin biosynthesis. Genistein affinity was more variable: GmIFH9A, GmIFH9C, GmIFH9F, and GmIFH15 displayed lower *K*_m_ values (4.88–11.85 µM), whereas GmIFH9D and GmIFH9E showed substantially higher *K*_m_ values (54.81 ± 6.20 and 65.20 ± 7.84 µM, respectively) Correspondingly, catalytic efficiencies (*k_cat_/K_m_*) for these substrates were highest among all reactions measured, particularly for GmIFH9A and GmIFH9E (Table 3).

**Table 3.** Michaelis-Menten kinetic parameters of GmI2’Hs with isoflavone substrates.

| I2'H | Substrates | V<sub>max</sub><br>(nmol/min/mg) | K<sub>m</sub> (μM) | k<sub>cat</sub> (s<sup>-1</sup>) | Catalytic<br>efficiency<br>(μM<sup>-1</sup>min<sup>-1</sup>) |
| --- | --- | --- | --- | --- | --- |
| GmIFH9A | Daidzein | 50.35 ± 1.53 | 7.61 ± 0.62 | 0.64 ± 0.02 | 5.08 |
|  | Genistein | 33.96 ± 1.88 | 4.88 ± 0.80 | 0.43 ± 0.02 | 5.34 |
|  | Glycitein | 12.96 ± 1.03 | 17.05 ± 3.16 | 0.17 ± 0.01 | 0.58 |
|  | Formononetin | 9.04 ± 0.28 | 14.12 ± 1.07 | 0.12 ± 0.00 | 0.49 |
|  | Biochanin A | 6.88 ± 0.36 | 93.59 ± 8.17 | 0.09 ± 0.00 | 0.06 |
| GmIFH9C | Daidzein | 3.11 ± 0.17 | 3.36 ± 0.69 | 0.04 ± 0.00 | 0.66 |
|  | Genistein | 8.10 ± 0.28 | 11.84 ± 0.98 | 0.10 ± 0.00 | 0.49 |
|  | Glycitein | 1.33 ± 0.13 | 3.83 ± 3.73 | 0.02 ± 0.00 | 0.04 |
| GmIFH9D | Daidzein | 1.62 ± 0.08 | 2.61 ± 0.55 | ND | ND |
|  | Genistein | 11.61 ± 0.71 | 54.81 ± 6.20 | ND | ND |
|  | Glycitein | 0.51 ± 0.03 | 7.49 ± 1.18 | ND | ND |
| GmIFH9E | Daidzein | 4.60 ± 0.27 | 4.65 ± 0.92 | 0.11 ± 0.01 | 1.37 |
|  | Genistein | 29.47 ± 1.98 | 65.20 ± 7.84 | 0.68 ± 0.05 | 0.62 |
|  | Glycitein | 1.58 ± 0.05 | 16.91 ± 1.20 | 0.04 ± 0.00 | 0.13 |
| GmIFH9F | Daidzein | 2.93 ± 0.17 | 4.28 ± 0.79 | 0.07 ± 0.00 | 1.01 |
|  | Genistein | 8.01 ± 0.35 | 11.85 ± 1.13 | 0.20 ± 0.01 | 1.00 |
|  | Glycitein | 0.50 ± 0.04 | 13.17 ± 2.00 | 0.01 ± 0.00 | 0.06 |
| GmIFH15 | Daidzein | 0.83 ± 0.12 | 4.99 ± 1.68 | 0.02 ± 0.00 | 0.27 |
|  | Genistein | 1.66 ± 0.13 | 10.20 ± 1.67 | 0.04 ± 0.00 | 0.26 |
|  | Glycitein | 0.34 ± 0.04 | 50.01 ± 10.06 | 0.01 ± 0.00 | 0.01 |

V<sub>max</sub>, K<sub>m</sub> and k<sub>cat</sub> are expressed as mean values of three independent replicates ±SE
ND, not determined

In contrast, glycitein generally showed markedly lower catalytic efficiencies than daidzein and genistein across most enzymes, indicating reduced overall turnover despite measurable activity. Notably, among the GmIFHs examined, GmIFH9A displayed the highest catalytic efficiency toward glycitein (0.58 µM⁻¹ min⁻¹), whereas GmIFH9C, GmIFH9E, GmIFH9F, and GmIFH15 showed substantially lower efficiencies (0.04, 0.13, 0.06, and 0.01 µM⁻¹ min⁻¹, respectively). Although glycitein *K*_m_ values were not uniformly high across all enzymes, its lower catalytic efficiencies indicate that it is a less favored substrate overall (Table 3).

Among the enzymes tested, GmIFH9A was uniquely competent in catalyzing 2′-hydroxylation of formononetin, exhibiting measurable turnover with a *K_m_* of 14.12 ± 1.07 µM and a catalytic efficiency of 0.49 µM⁻¹ min⁻¹. Although the catalytic efficiency for formononetin was lower than that observed for daidzein and genistein with the same enzyme, it is consistent with the formation of 2′-hydroxyformononetin detected in enzymatic assays and soybean tissues (Table 3). For GmIFH9D, *k*cat and catalytic efficiency were not determined due to the lack of a suitable detectable peptide for LC-MS-based quantification; however, its *K*_m_ and Vmax values indicate stronger apparent affinity toward daidzein than genistein or glycitein.

### Metabolomics analysis of hydroxyisoflavones in soybean root tissue

To confirm that the newly discovered 2′-hydroxyglycitein and 2′-hydroxyformononetin occur in soybean roots, we used a targeted, high resolution liquid chromatography with mass spectrometry (LC-MS/MS) approach. Low levels of hydroxyisoflavones were detectable by MS/MS in soybean root extract samples; however, these were below the level of detection for confident identification by MS/MS due to their low abundance. Conducting acid hydrolysis of the samples resulted in the detection and identification of several hydroxyisoflavones in the soybean root tissue including the two newly found 2′-hydroxyglycitein and 2′-hydroxyformononetin (Figure 5). The accurate mass values and diagnostic MS/MS fragmentation patterns of the hydrolyzed plant extracts were identical to those obtained for enzymatically synthesized reference standards of the hydroxyisoflavones. This confirmed the structural identity of the hydroxyisoflavones and their presence *in planta*.

**Figure 5.**
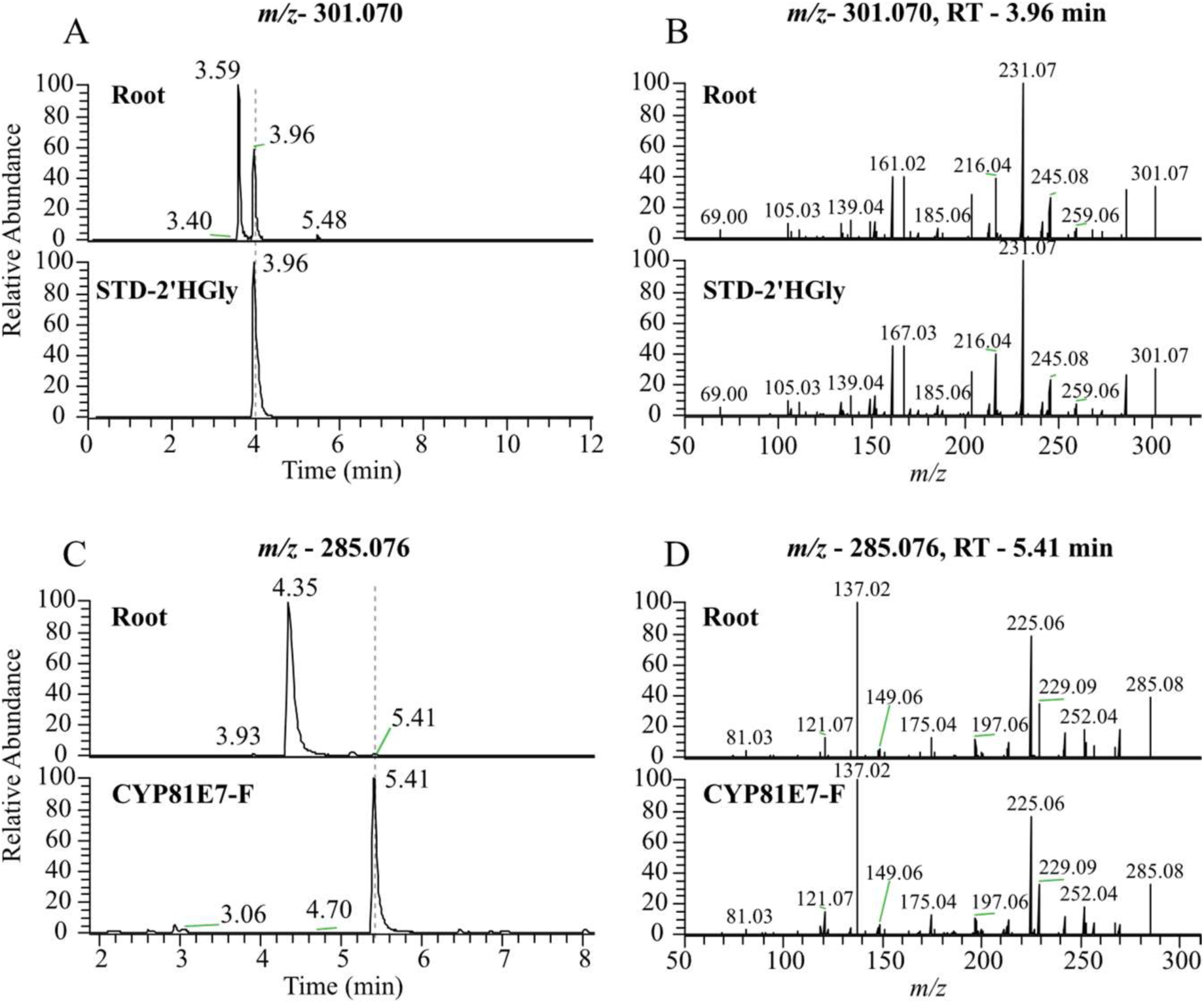
Confirmation of 2’-hydroxyglycitein and 2’-hydroxyformononetin in soybean roots. **(A)** Extracted base peak chromatograms at *m/z* 301.070 showing 2’-hydroxyglycitein (2’HGly) at retention time (RT) 3.96 min in soybean root extract and NMR-verified 2-′HGly reference compound purified from GmIFH enzyme assays (STD), **(B)** MS/MS fragmentation of the samples in A, **(C)** Base peak at *m/z* 285.076 showing 2’-hydroxyformononetin at retention time (RT) 5.41 min in soybean root extract and 2’-hydroxyformononetin product formed in positive control *M. truncatula* I2’H CYP81E7 enzyme assay incubated with formononetin as substrate, **(D)** MS/MS fragmentation of the samples in C.

## Discussion

### Expansion of soybean hydroxyisoflavone diversity and new phytoalexin branch points

Hydroxylated isoflavones are key metabolic branch points of legume specialized metabolism enabling redirection of central isoflavone aglycones to varied defense-related pathways. Canonical examples include 2′-hydroxydaidzein which take part in the production of glyceollins in soybean and as an intermediate linked to coumestrol formation in several legumes establishing a connection between stress perception and antimicrobial metabolite deployment (Jeon et al., 2012; Sukumaran *et al*., 2018). In *M. truncatula*, the 2′-hydroxyformononetin is important in the production of pterocarpan phytoalexin medicarpin, whilst 3′-hydroxyformononetin is linked with defence associated compounds like maackiain and pisatin in pea, further supporting the role played by hydroxylation regiochemistry in pathway commitment (Hinderer et al., 1987; Liu *et al*., 2003). In a parallel manner, 2′-hydroxygenistein has been suggested to be an intermediate in the kievitone biosynthesis of *Phaseolus* species (O’Neill et al., 1983). Unlike these well-characterized intermediates, 2′-hydroxyglycitein was not reported before.

Recent studies have also established a close association between glycitein biosynthesis and genetic regulation of *P. sojae* resistance in soybean, indicating that glycitein is a functional defense node and not an isoflavone peripheral pool (Yang et al., 2025). This study provides the direct evidence for presence of 2′-hydroxyglycitein in soybean roots with the aid of an enzymatically produced reference standard along with the targeted LC-MS/MS (Figure 6). Even though the downstream fate of 2′-hydroxyglycitein is yet to be determined, legume hydroxyisoflavone precedents suggest that these intermediates are often further tailored through prenylation, methylation, and/or cyclization to produce bioactive defense metabolites (Araya-Cloutier et al., 2017; Levisson et al., 2019; Shen et al., 2012). Soybean accumulates a diverse array of prenylated isoflavones following microbial and chemical elicitation, which gives a viable biochemical situation where 2′-hydroxyglycitein could act as an initial step towards further specialized-metabolic products (Kalli et al., 2020). Simultaneously, the presence of 2′-hydroxyformononetin in soybean roots (Figure 5) suggests the existence of inducible isoflavonoid branches beyond the canonical daidzein-centered glyceollin pathway. In *Medicago*, a major intermediate in the medicarpin pathway is 2′-hydroxyformonetin (Liu *et al*., 2003). Medicarpin detected in recent metabolomics experiments in soybean further support the idea that soybean can use other routes involving pterocarpan-type phytoalexin under specific contexts (de Souza Gouveia et al., 2025; Zhao et al., 2025). Consistent with this idea, an o-methylated isoflavone feature, which was observed in our previous dataset (*m/z* 299.091, RT 3.33 min), may be a possible downstream methylation of 2′-hydroxyformononetin by an o-methyltransferase (Anguraj Vadivel *et al*., 2021) (Figure 6). Together, these discoveries of 2′-hydroxyglycitein and 2′-hydroxyformononetin demonstrate the importance of previously overlooked metabolic intersections in soybean and highlight the hidden flexibility within its stress-responsive isoflavonoid network.

**Figure 6.**
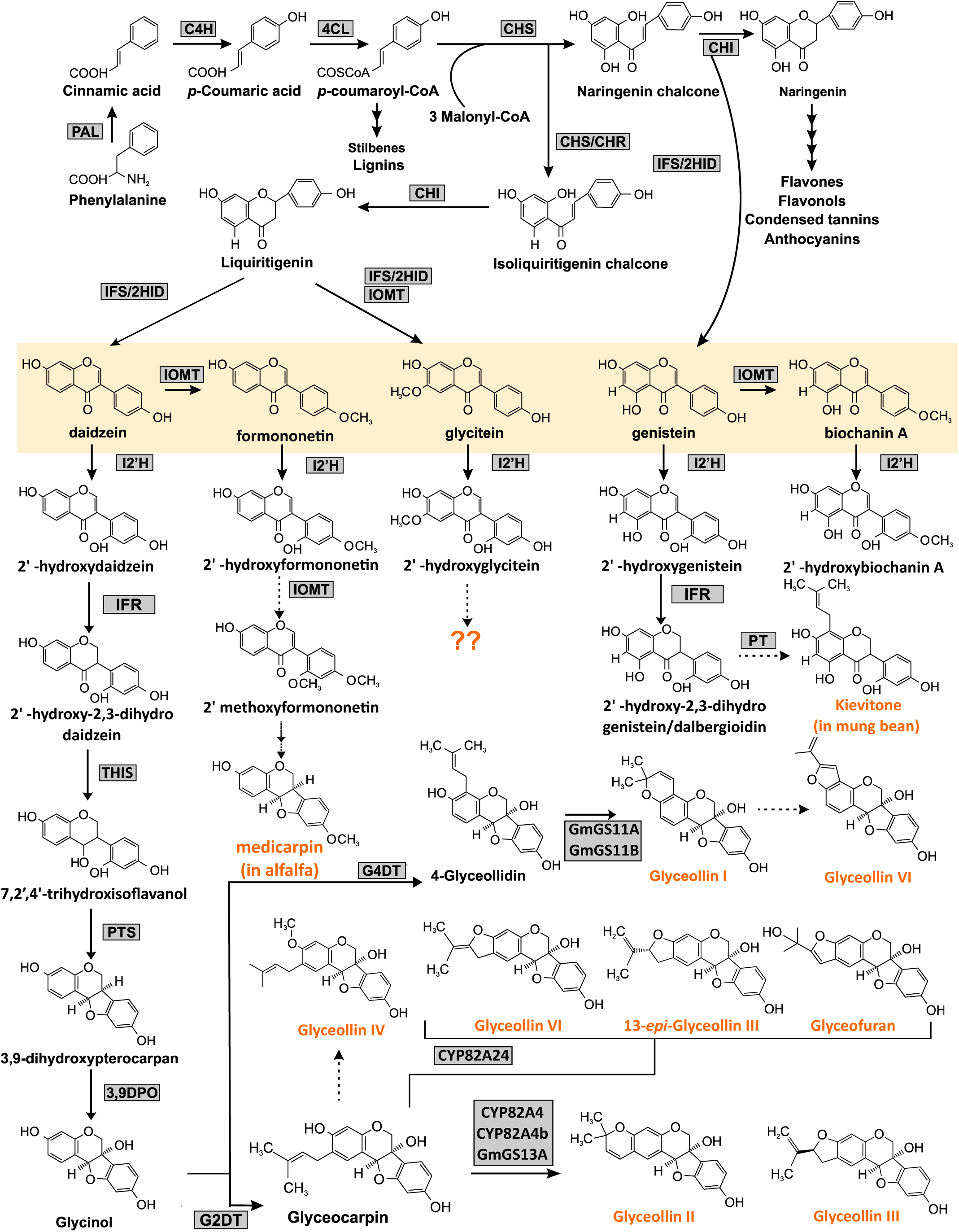
Isoflavonoid biosynthetic pathway in soybean showing proposed downstream products from hydroxyisoflavones. Isoflavone aglycons are highlighted (yellow box). Compounds in orange text represent putative isoflavones derived from different 2’-hydroxyisoflavones. PAL, phenylalanine ammonia-lyase; C4H, cinnamate 4-hydroxylase; 4CL, 4-coumarate:CoA ligase; CHS, chalcone synthase; CHR, chalcone reductase; CHI, chalcone isomerase; IFS, isoflavone synthase; 2HID, 2-hydroxyisoflavanone dehydratase; I2′H, isoflavone 2′-hydroxylase; IFR, isoflavone reductase; THIS, 7,2′,4′-trihydroxyisoflavanol synthase; PTS, pterocarpan synthase; 3,9DPO, 3,9-dihydroxypterocarpan 6a-hydroxylase; G4DT, glycinol 4-dimethylallyltransferase; G2DT, glycinol 2-dimethylallyltransferase; IOMT, isoflavone O-methyltransferase. Dotted arrows show predicted steps, whereas multiple arrows represent multiple steps.

### IFH family expansion as a mechanistic basis for hidden hydroxylated intermediates

Soybean encodes an expanded CYP81E repertoire in the context of a highly duplicated paleopolyploid genome, giving a vast possibility of retention and divergence of paralogous pathway enzymes (Schmutz et al., 2010). Such duplication-driven diversification has been identified as one way that plant specialized metabolism can acquire new branch capacity, by sub-and neo-functionalization, and is often produced by the promiscuous ancestral activities coupled with subsequent refinement (Vogt, 2010). Our *in vitro* enzyme assays indicate that several soybean IFHs are catalytically active and comprehensively span both 2′- and 3′-hydroxylation chemistries, though individual IFHs vary significantly in substrate specificity (Figure 4). This extended and diversified IFH ensemble offers a mechanistic explanation of the appearance of these hidden hydroxylated intermediates which can either be cryptic due to low flux or may be accumulated under certain stress conditions. The stress inducible nature of glycitein-derived branches under biotic stress in soybean further supports the argument (Uchida et al., 2020) that a permissive enzyme (e.g., GmIFH9A) that is capable of producing hydroxylated products of many different aglycones can create alternative hydroxylated entry points (including those derived from glycitein and formononetin), while other paralogs contribute more specialized, canonical activities. Together, soybean IFH family expansion and functional divergence is a direct mechanism to explain why such hydroxylated intermediates such as 2′-hydroxyglycitein and 2′-hydroxyformononetin might be present as unresolved components of the soybean defense metabolome.

### GmIFH9A as a permissive catalyst connecting glyceollin-associated metabolism with parallel branches

Among the soybean IFH isoforms, GmIFH9A is notable due to its remarkably broad substrate range and dual 2′/3′ regioselectivity, transforming all the five isoflavone aglycones and being the only enzyme with strong activity with both formononetin and biochanin A (Figure 4). This biochemical profile is especially pertinent to the metabolite-driven story discovered in this study since it offers a direct enzymatic pathway to both 2′-hydroxyglycitein and 2′-hydroxyformononetin intermediates that emerged as proposed by feature mining in the GmMYB176-GmbZIP5 metabolomics (Anguraj Vadivel *et al*., 2021) dataset and are presently validated by our *in planta* finding. From an evolutionary and pathway-logic perspective, one of the possibilities is that a permissive IFH like GmIFH9A is a catalytic scaffold which gave rise to more specialized soybean IFHs by subsequent duplication, followed by a restriction of substrate tolerance. Under this model, the apparent loss of activity toward 4′-methoxy isoflavones in several soybean I2′Hs would reflect specialization toward the dominant soybean aglycone pool (e.g., daidzein/genistein), whereas retention of glycitein and formononetin competence in select enzymes would preserve the capacity to seed alternative hydroxylated branches when conditions favor them. A similar contrast between permissive and restricted catalytic behavior has already been documented for other soybean isoflavonoid P450s, where closely related CYP93A paralogs differ in substrate breadth and their altered expression measurably shifts isoflavonoid flux in planta (Xia et al., 2023). This interpretation aligns with broader general specialised-metabolic evolutionary theories which postulate that enzyme promiscuity may be pre-pathway diversification and may facilitate the formation of cryptic branches only to be revealed under specific transcriptional or elicitation conditions (Moghe and Last, 2015).

### Physiological relevance and downstream implications of newly validated hydroxylated branch points

The central advance of this study is not solely the biochemical potential of soybean IFHs *in vitro*, but in directly connecting those activities to authentic soybean metabolites by using enzymatically generated standards and targeted MS/MS validation in soybean roots. The detection of 2′-hydroxyglycitein in soybean roots confirms that this modification is an intrinsic feature of the soybean metabolome, rather than an artifact limited to heterologous expression systems. On the same note, the presence of 2′-hydroxyformononetin supports the presence of inducible metabolic pathways that do not rely on the canonical glyceollin axis, as is consistent with emerging evidence that soybean can accumulate additional pterocarpan-like defensive metabolites under particular conditions (de Souza Gouveia *et al*., 2025; Zhao *et al*., 2025). Notably, these observations improve the existing pathway models by implying that defense metabolism of soybean can be better described as a network of conditionally interconnected branches and not as a single glyceollin pipeline. In practice, the feasibility of catalytically efficient soybean IFHs especially of enzymes that generate hydroxylated methoxy-isoflavones can offer opportunities to (i) make synthetic standards for resolving the unknown metabolomics features, (ii) probe downstream tailoring reactions (prenylation, O-methylation, cyclization) that could convert these intermediates into bioactive end products, and (iii) provide opportunities to scale with hydroxyisoflavones and their derivatives using microbial or plant engineering to test their roles in plant resilience. Together, our findings reveal two previously unknown hydroxylated compounds 2′-hydroxyglycitein and 2′-hydroxyformononetin which broaden the chemical space of soybean isoflavonoids and form a mechanistic foundation of other, concealed defense pathways in addition to glyceollin.

## Material and methods

### Plant material, analytical standards and chemicals

The seeds of soybean (*Glycine max* [L] Merr.) cultivar Williams 82 were germinated in sterile vermiculite at 27 °C under 16 h with 450 µmoles m^-2^ s^-1^ light and 18 °C for 8 h at London Research and Development Centre. Soybean seedlings were infected with *Phytophthora sojae* zoospores as described previously (Khatri et al., 2022). Root tissues were harvested 24 h post-inoculation, flash frozen in liquid N_2_ and stored at -80 °C until needed.

The HPLC grade standards of daidzein, genistein and glycitein were obtained from LC laboratories (Woburn, MA). Formononetin, biochanin A and 3′-hydroxydaidzein were purchased from Cayman Chemicals (Ann Arbor, MI). 2′-hydroxybiochanin A was obtained from Biosynth (Compton, UK). 3′-hydroxygenistein (Orobol), 3′-hydroxyformononetin (Calycosin), and 3′-hydroxybiochanin A (Pratensein) were obtained from (MedChemExpress, NJ). For HPLC analysis, analytical-grade chemicals and solvents were obtained from Sigma-Aldrich and Thermo-Scientific, Canada.

### Isoflavone-related metabolite features mining

The untargeted metabolomics dataset (Accession #ST001634) that contains LC-MS data from the GmMYB176-GmbZIP5 overexpressing soybean hairy roots (Anguraj Vadivel et al., 2021) was re-analyzed through the Metabolomics Workbench platform (https://www.metabolomicsworkbench.org). Theoretically *m/z* values corresponding to the five major legume isoflavones with hydroxylated, glycosylated, and malonylated derivatives were calculated and used to interrogate the feature list using a mass tolerance of ±5 ppm. From this putative list, features exhibiting ≥2-fold accumulation and FDR value < 0.05 in overexpressed samples relative to control samples were retained for further analysis (Supplementary Table 1). The volcano plot was generated using *ggplot* (Wilkinson, 2011) in R studio and self-organizing map was constructed following the methodology described previously (Dang et al., 2018) using *kohonen* (Kohonen, 1998) package. The retention times and MS/MS spectra of these candidate features were compared with enzymatically produced reference standards generated in this study to enable structural assignment.

### Identification of CYP81E candidates, phylogenetic and motif analysis

CYP81E candidates were identified based on our previously catalogued CYP81 in soybean (Khatri *et al*., 2022). The candidates were further validated based on their sequence homology and clustering with functionally characterized IFHs from legumes including *M. truncatula* I2′H (CYP81E7; Uniprot: Q6WNR0) and I3′H (CYP81E9; Uniprot: Q6WNQ9), *L. japonicus* I2′H (Uniprot: Q9MBE4) and *G. echinata* I2′H (CYP81E1; Uniprot: P93147). Additionally, amino acid sequences of characterized CYP81 members from other plant species were retrieved from literature search (Supplementary Table 2). Multiple sequence alignment was generated using Clustal Omega (Sievers et al., 2011) and phylogenetic tree was generated using the maximum-likelihood method implemented in IQ-TREE with automatic best-fitting evolutionary model selection (http://iqtree.cibiv.univie.ac.at/) (Trifinopoulos et al., 2016). The JTT+R4 substitution model was selected, and node support was evaluated using 1000 ultrafast bootstrap replicates. CYP102A1 was used as an outgroup to root the phylogenetic trees and visualized using MEGA 12 (Kumar et al., 2024), Predicted molecular weight and isoelectric point (pI) values of GmIFH proteins were calculated using the ExPASy ProtParam server (https://web.expasy.org/compute_pi/).

Conserved cytochrome P450 signature motifs were identified by manual inspection of protein sequences aligned using Clustal omega and visualized using pyBoxshade (https://github.com/mdbaron42/pyBoxshade). Pairwise amino acid sequence identity among GmIFHs was calculated using SIAS tool (http://imed.med.ucm.es/Tools/sias.html). The analysis was conducted using BLOSUM62 matrix, with gap penalties set at a creation cost of 10 and an extension cost of 0.5.

### Gene cloning, heterologous expression, and microsome preparation

Full length coding sequences of *GmIFH* candidates were amplified from soybean cDNA using gene-specific primers (Supplementary Table 3) and recombined into the Gateway-compatible pDONR-Zeo entry vector (Earley et al., 2006) and then into pESC-Leu2d-LjCPR1-GW destination vector (Khatri et al., 2023) and were transformed into *Saccharomyces cerevisiae* strain BY4742. Yeast cultures were induced, and microsomes containing recombinant GmIFH were prepared by differential centrifugation as previously described (Khatri *et al*., 2023). Microsomal protein concentrations were determined using the Bradford assay.

### MS-based proteomic quantification of recombinant GmIFHs in yeast microsomes

Microsomal protein (75 µg) was denatured in 400 µL digestion buffer containing 1% sodium deoxycholate (SDC) and 100 mM Tris-HCl (pH 8.5) and incubated at 60 °C for 10 min. Proteins were reduced with freshly prepared DTT (final concentration 1.9 mM) at 60 °C for 1 h and alkylated with iodoacetamide (final concentration 4.3 mM) for 30 min in the dark at room temperature. Excess iodoacetamide was quenched by additional DTT treatment for 5 min. Proteolytic digestion was performed by adding trypsin (enzyme-to-protein ratio 1:10, w/w) and incubating at 37 °C for 24 h. Following digestion, SDC was precipitated by the addition of 1% trifluoroacetic acid, removed from the tryptic peptides by ethyl acetate, liquid-liquid extraction. The remaining aqueous phase containing the peptides was diluted with Milli-Q water and desalted using Oasis HLB solid-phase extraction cartridges 30 mg sorbent bed (Waters Corporation, Milford, CT). Cartridges were activated with methanol, equilibrated with 0.1% formic acid. After sample loading and drying, peptides were eluted by two additions of 200 µL of 70% acetonitrile. Eluates were dried under nitrogen gas and reconstituted in 200 µL 5% acetonitrile containing 0.1% formic acid prior to LC–MS/MS analysis. The concentration of the expressed enzymes within the total microsomal protein tryptic digest were determined by synthesizing sequence-specific peptides of GmIFH enzymes (GenScript, USA) (Supplementary Table 4) and using those to build a calibration curve. The standards along with the microsomal protein tryptic digest were analyzed on a Thermo Altis triple quadrupole mass spectrometer to estimate the abundance of recombinant GmIFHs in the microsomal fraction.

### *In vitro* enzyme assay and kinetic analysis

*In vitro* enzyme assays with yeast microsomal proteins containing recombinant GmIFH was performed as described in Khatri *et al*., (2023). Microsomal proteins (0.5 mg) were incubated with 50 mM phosphate buffer (pH 7.6) containing 1 mM NADPH and 20 µM of each of the isoflavone aglycones (daidzein, genistein, glycitein, formononetin and biochanin A) in an independent reaction at 25 °C for 5 hours. The reaction was stopped with 10 µL of concentrated HCl, followed by two ethyl acetate extractions. The extract was evaporated under N_2_ gas and residue was resuspended in 100 µL methanol for subsequent HPLC analysis.

To determine apparent *K*_m_ values, GmIFH microsomal protein (2 to 500 µg) was incubated with substrate concentrations ranging from 1 to 120 µM for 15 to 120 mins. Other reaction components were kept the same as described above, followed by HPLC analysis. Substrate depletion method was adopted for kinetic study. Kinetic constants were calculated from initial rate data using the Michaelis Menten equation in GraphPad Prism 8 using nonlinear regression. The reaction substrate was compared with the authentic standards.

### Whole yeast feed assay, purification of enzyme products and NMR analysis

To generate sufficient quantities of GmIFH reaction products for structural elucidation, yeast clones expressing GmIFH9A were cultured in selective dropout medium as described above. Following growth in glucose-containing medium, cells were harvested by centrifugation and resuspended in induction medium. Individual isoflavone substrates (daidzein, genistein, or glycitein) were added to separate cultures at a final concentration of 30 µM, and incubated for 24 h. Cultures were centrifuged at 10,000 × *g*, and supernatants were collected and extracted twice with an equal volume of ethyl acetate. Combined organic extracts were concentrated using a Buchi R-300 (New Castle, DE) rotary evaporator under reduced pressure (240 mbar) to a final volume of 5-10 mL, followed by complete solvent removal under nitrogen gas. Residues were reconstituted in 2 mL methanol and subjected to semi-preparative reverse-phase HPLC using a Gemini C18 column (10 × 150 mm, 5 µm; Phenomenex, Torrence, CA) at a flow rate of 3 mL min⁻¹, employing previously described chromatographic conditions. Target product peaks were collected using an Agilent G1364C fraction collector coupled to an Agilent 1260 HPLC system. Collected fractions were lyophilized using a freeze dryer (Labconco, KS), verified for product identity, and subsequently used for NMR-based structural characterization. NMR spectra for structural elucidation were acquired in dimethyl sulfoxide (DMSO-d_6_) using a high-field Bruker Neo 600 MHz spectrometer equipped with an H/FX 5 mm Bruker iprobe at the JB Stothers NMR Facility, Western University.

### Metabolite extraction and acid hydrolysis

Metabolite extraction from soybean root tissues was performed using lyophilized samples (10 mg) in 1 mL 80% methanol (v/v). The samples were sonicated in an ice water bath for 20 min, followed by centrifugation at 11,000 × g for 10 min at ambient temperature. The supernatant (700 µL) was dried under nitrogen gas then redissolved in 200 µL of 50% methanol and filtered through a 0.45 µm syringe filter (Millipore, U.S.). Acid hydrolysis of the samples was performed on soybean root extracts following the previously established method (Delmonte et al., 2006).

### HPLC and LC-MS analysis

HPLC analysis was performed following methods previously developed using an Agilent 1260 (Santa Clara, CA) series HPLC (Khatri *et al*., 2023) with the following modifications. A Kinetex XB-C18 (Phenomenex, Torrence, CA) column was used for the analysis of daidzein, genistein and glycitein, and separation occurred at a flow rate of 0.8 mL min⁻¹. Spectra were monitored at 254 nm with a photodiode array detector with an injection volume of 10 µL. The gradient of solvent B was as follows: 12% from 0 to 1 min, 12–35% from 1 to 16 min, 80% from 16 to 17 min, 100% from 17.5 to 18 min, followed by re-equilibration at 12% until 20 min. For formononetin and biochanin A the gradient elution of solvent B was as follows: 35% from 0 to 1 min, 35–75% from 1 to 10 min, 100% from 10 to 12.5 min, followed by re-equilibration at 35% for 2 min. The UV spectrum and retention time of isoflavone substrates were compared to commercial standards while products were confirmed with NMR. CYP81E7 (Liu *et al*., 2003) was also used as a positive control.

For the analysis of isoflavones and hydroxyisoflavones in soybean tissue, a high-resolution Q-Exactive Quadrupole Orbitrap mass spectrometer (Thermo Fisher Scientific, U.S.) coupled with an Agilent 1290 HPLC system was used. The HPLC system was configured with a Zorbax Eclipse Plus RRHD C18 column (2.1 × 50 mm, 1.8 μm), maintained at a temperature of 35 °C. Samples of 5 μL each were injected at a flow rate of 0.3 mL min⁻¹. Mobile phases A and B, consisting of water with 0.1% formic acid and acetonitrile with 0.1% formic acid, respectively (Optima grade, Fisher Scientific, U.S.), were utilized. Mobile phase B was initiated at 0% for 0.5 min, followed by a gradual increase to 100% over 3.0 min. It was maintained at 100% for 2 min before reverting to 0% over 30s. Heated electrospray ionization (HESI) conditions included a spray voltage of 3.9 kV (HESI+), capillary temperature at 400 °C; probe heater temperature at 450 °C; sheath gas at 17 arbitrary units; auxiliary gas at 8 arbitrary units; and S-Lens RF level at 45. Compound detection occurred at a resolution of 140,000, employing full mass scans within the *m/z* range of 100 to 1200 in separate positive and negative ionization runs. Automatic gain control (AGC) target and maximum injection time (IT) were set at 3 × 10⁶ and 512 ms, respectively.

For structural confirmation, selected hydroxyisoflavone precursor ions were subjected to targeted MS/MS analysis using a 1.2 *m/z* isolation window, normalized collision energy (NCE) of 50, and exclusion from subsequent MS/MS for 5 sec. Compounds were identified using enzyme assay products and available commercial standards. Raw files were analyzed using Xcalibur Qual Browser (Thermo Fisher, USA)

### Conclusions

Our findings reposition soybean isoflavonoid metabolism as a more expansive and flexible network than previously described, in which hydroxylation serves not only to initiate glyceollin biosynthesis but also to seed additional, conditionally activated defense branches. By integrating metabolomics-guided feature mining with enzymology and *in planta* validation, we identify 2′-hydroxyglycitein and 2′-hydroxyformononetin as previously hidden hydroxylated entry points that extend the chemical space of soybean specialized metabolism. The enzymatic accessibility of these intermediates particularly through permissive IFHs such as GmIFH9A provides a mechanistic basis for their formation and offers tools to interrogate their downstream fates. More broadly, this work illustrates how latent pathway capacity encoded within expanded enzyme families can be revealed through stress-responsive regulation, yielding metabolic diversity that may contribute to plant resilience.

## Acknowledgements

We thank Kuflom Kuflu, and Alex Molnar (AAFC-London) for technical assistance and Dr. Mathew Willans at the JB Stothers NMR facility at Western University for help with NMR analysis. This research was supported by Agriculture and Agri-Food Canada’s Genomics Research and Development Initiative grant (J-002364) and the Natural Sciences and Engineering Research Council of Canada (NSERC) Discovery Grant 044661-2018 RGPIN to SD. PK was also supported by Queen Elizabeth II Graduate Scholarship in Science and Technology.

---

**Supplementary Figure 1:** Self-organizing map (SOM) outputs for metabolite clustering analysis. The SOM was created based on the metabolite abundance data of control (Ctrl_1-Ctrl_5) and *GmMYB176-GmbZIP5* overexpression (OE_1-OE_5) soybean hairy-root samples (Anguraj Vadivel et al., 2021). **(A)** Number of metabolite features per SOM node, showing the distribution of features across the 10 × 10 map, **(B)** Neighbour distances (U-matrix): Euclidean distances of adjacent nodes, with warmer colours denoting a closer relationship and lighter colours denoting a stronger separation between neighbouring clusters, **(C)** SOM codebook vectors: representative abundance patterns of each node, displaying the relative feature profiles of all OE and control samples. All the metabolites that are assigned to each node are summarized as the average pattern and the green color of the plot represents the samples of OE and the beige/ gray color of the plot represents the samples of the control.

**Supplementary Figure 2.** Multiple sequence alignment of deduced amino acid sequences of candidate GmIFHs with characterized I2’H (CYP81E1, CYP81E7, CYP81E63, LjCYP-2) and I3’H (CYP81E9) from other legumes. Identical and similar amino acid residues are indicated by black and gray shades, respectively. Five P450 motifs: the PPGP (pink), the oxygen-binding (green), the ETLR (blue), the PERF (yellow) and the heme-binding motifs (purple) are indicated by different colored rectangles. Only an abridged version of the alignment is shown.

**Supplementary Figure 3.** GmIFH sequence identity. Sequence identity heatmap showing the pairwise percentage identity between GmIFH amino acid sequences. The X and Y axes indicate 11 identified GmIFHs. Identity scores are shown as a color-coded matrix, calculated by comparing every sequence to each other. Scale bar showing increase in sequence identity from white to green. Percent identity is also shown in each box.

**Supplementary Figure 4.** Functional characterization of candidate GmIFHs *in vitro*. HPLC chromatogram showing the substrate and product peaks detected at 254 nm UV detector. Microsomal fraction containing candidate GmIFHs and LjCPR1 was incubated with different isoflavone aglycone substrates and NADPH followed by ethyl acetate extraction and analyzed by HPLC. Dotted lines indicate substrate and product peaks. 2’-OHD: 2’-hydroxydaidzein, 3’-OHD: 3’-hydroxydaidzein, 2’-OHG: 2’-hydroxygenistein, 3’-OHG: 3’-hydroxygenistein, 2’-OHGly: 2’-hydroxyglycitein, 3’-OHGly: 3’-hydroxyglycitein, F: Formononetin, 2’-OHF: 2’-hydroxyformononetin, 3’-OHF: 3’-hydroxyformononetin, B: Biochanin A, 2’-OHB: 2’-hydroxybiochanin A, 3’-OHB: 3’-hydroxybiochanin A. CYP81E7: *Medicago truncatula* I2’H.

**Supplementary Figure 5:** ^1^H and ^13^C NMR spectra of 2’-hydroxyglycitein in DMSO-*d_6_*.

**Supplementary Figure 6:** COSY and HSQC NMR spectra of 2’-hydroxyglycitein in DMSO-*d_6_*.

**Supplementary Figure 7:** HMBC NMR spectra of 2’-hydroxyglycitein in DMSO-*d_6_*.

**Supplementary Figure 8:** ^1^H and ^13^C NMR spectra of 2’-hydroxydaidzein in DMSO-*d_6_*.

**Supplementary Figure 9:** COSY and HSQC NMR spectra of 2’-hydroxydaidzein in DMSO-*d_6_*.

**Supplementary Figure 10:** HMBC NMR spectra of 2’-hydroxydaidzein in DMSO-*d_6_*.

**Supplementary Figure 11:** ^1^H and ^13^C NMR spectra of 2’-hydroxygenistein in DMSO-*d_6_*.

**Supplementary Figure 12:** COSY and HSQC NMR spectra of 2’-hydroxygenistein in DMSO-*d_6_*.

**Supplementary Figure 13:** HMBC NMR spectra of 2’-hydroxygenistein in DMSO-*d_6_*.

**Supplementary Figure 14.** MS spectra showing the presence of hydroxyglycitein and hydroxyformononetin in the GmIFH enzyme assay **(A)** Base peak at *m/z* 301.070 showing 2’-hydroxyglycitein (2’HGly) and 3’-hydroxyglycitein (3’HGly) at retention times (RT) of 3.96 and 3.78 min, respectively, in 2’HGly NMR verified compound standard (STD) and GmIFH9A, GmIFH9B and *Medicago truncatula* I2’H (CYP81E7) enzyme assay reactions with substrate glycitein (Gly). **(B, C)** MS/MS fragmentation of samples in panels A. **(D)** Base peak at *m/z* 285.075 showing 2’-hydroxyformononetin (2’HF) and 3’-hydroxyformononetin (3’HF) at retention times (RT) of 5.41 and 4.52 min, respectively, in GmIFH9A and GmIFH8A enzyme assay reactions incubated with substrate formononetin (F). *Medicago truncatula* I2’H (CYP81E7) enzyme assay reaction with substrate F used as a positive control for confirming 2’HF. **(E, F)** MS/MS fragmentation of samples in panel D.

**Supplementary Figure 15.** MS spectra showing the presence of hydroxydaidzein and hydroxygenistein in the GmIFH enzyme assay. **(A)** Base peak at *m/z* 271.060 showing 2’-hydroxydaidzein (2’HD) and 3’-hydroxydaidzein (3’HD) at retention times (RT) of 3.81 and 3.67 min, respectively, in 2’HD, 3’HD standards and GmIFH9A and GmIFH9B enzyme assay reactions incubated with substrate daidzein (D), **(B, C)** MS/MS fragmentation of samples in panels A, **(D)** Base peak at *m/z* 287.055 showing 2’-hydroxygenistein (2’HG) and 3’-hydroxygenistein (3’HG) at retention times (RT) of 4.39 and 4.48 min, respectively, in 2’HG NMR verified compound standard (STD) and GmIFH9A and GmIFH9B enzyme assay reactions incubated with substrate genistein (G) The *Medicago. truncatula* I2′H/CYP81E7 enzyme assay with genistein was used as a positive control for 2′-hydroxygenistein identification, **(E, F)** MS/MS fragmentation of samples in panel D.

**Supplementary Figure 16.** MS spectra showing the presence of hydroxybiochanin A in the GmIFH enzyme assay. **(A)** Base peak at *m/z* 301.070 showing 2’-hydroxybiochanin A (2’HB) and 3’- hydroxybiochanin A (3’HB) at retention times (RT) of 5.79 and 5.51 min, respectively, in 2’HB standard and GmIFH9A and GmIFH8A enzyme assay reactions with substrate biochanin A (B). *Medicago truncatula* I2’H (CYP81E7) enzyme assay reaction with substrate B used as a positive control for confirming 2’HB. **(B, C)** MS/MS fragmentation of samples in panels A.

**Supplementary Figure 17.** Confirmation of hydroxydaidzein and hydroxygenistein in soybean roots. **(A)** Base peak at *m/z* 271.060 showing 2’-hydroxydaidzein (2’HD) and 3’-hydroxydaidzein (3’HD) at retention times (RT) of 3.71 and 3.57 min, respectively, in soybean root extract and authentic standards of 2’HD and 3’HD. **(B, C)** MS/MS fragmentation of samples in panels A, **(D)** Base peak at *m/z* 287.055 showing 2’-hydroxygenistein (2’HG) at RT 4.30 min in soybean root extract and the 2’HG NMR verified compound standard, **(E)** MS/MS fragmentation of the sample in panel D.

