## Supplementary Figures for "Multi-Substrate Specificity of Isoflavone hydroxylases (GmIFH) Drive Isoflavonoid Diversification in Soybean"

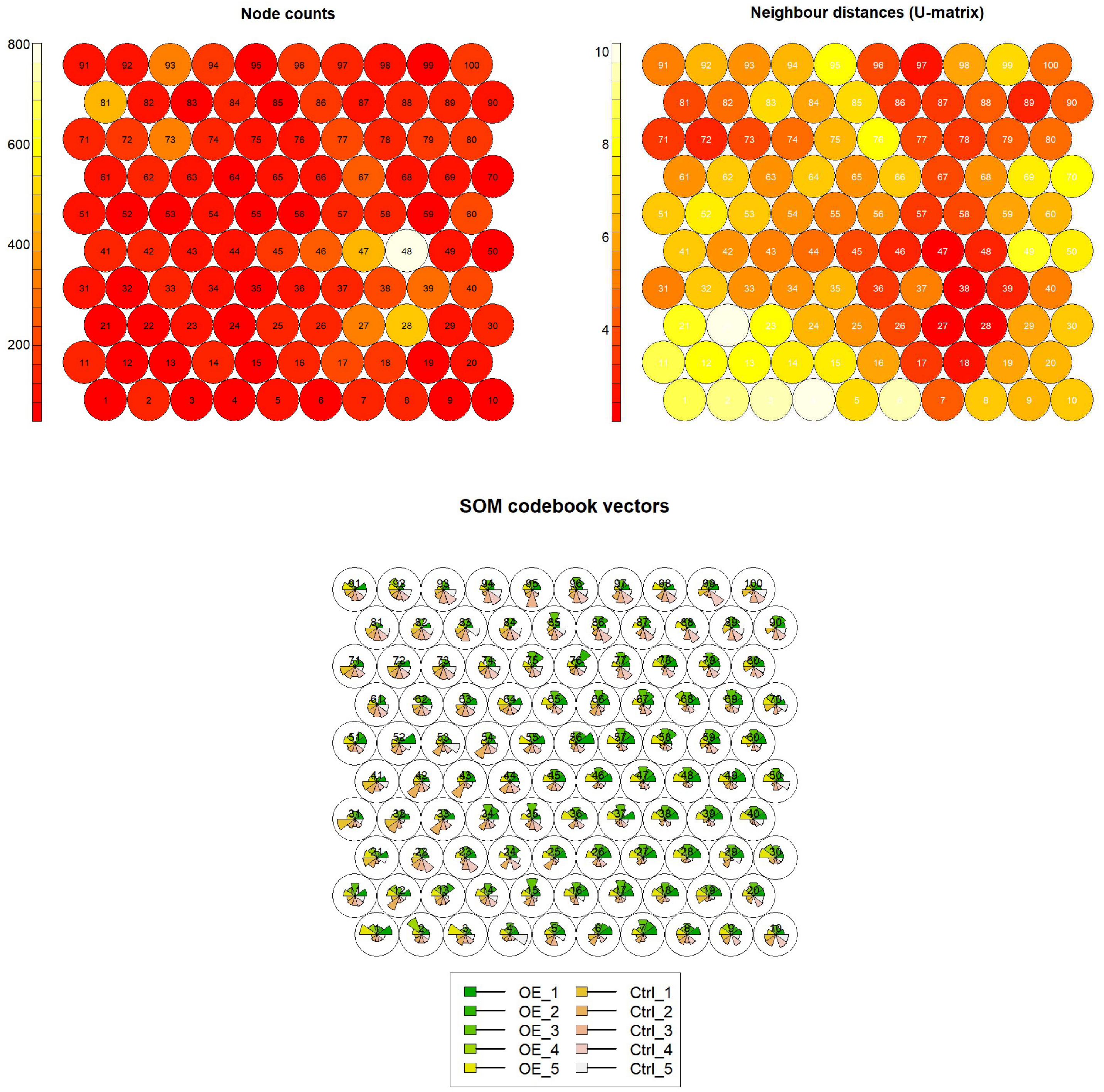


**Supplementary Figure 1:** Self-organizing map (SOM) outputs for metabolite clustering analysis. The SOM was created based on the metabolite abundance data of control (Ctrl_1-Ctrl_5) and *GmMYB176-GmbZIP5* overexpression (OE_1-OE_5) soybean hairy-root samples (Anguraj Vadivel et al., 2021). (A) Number of metabolite features per SOM node, showing the distribution of features across the 10 × 10 map, (B) Neighbour distances (U-matrix): Euclidean distances of adjacent nodes, with warmer colours denoting a closer relationship and lighter colours denoting a stronger separation between neighbouring clusters, (C) SOM codebook vectors: representative abundance patterns of each node, displaying the relative feature profiles of all OE and control samples. All the metabolites that are assigned to each node are summarized as the average pattern and the green color of the plot represents the samples of OE and the beige/ gray color of the plot represents the samples of the control.

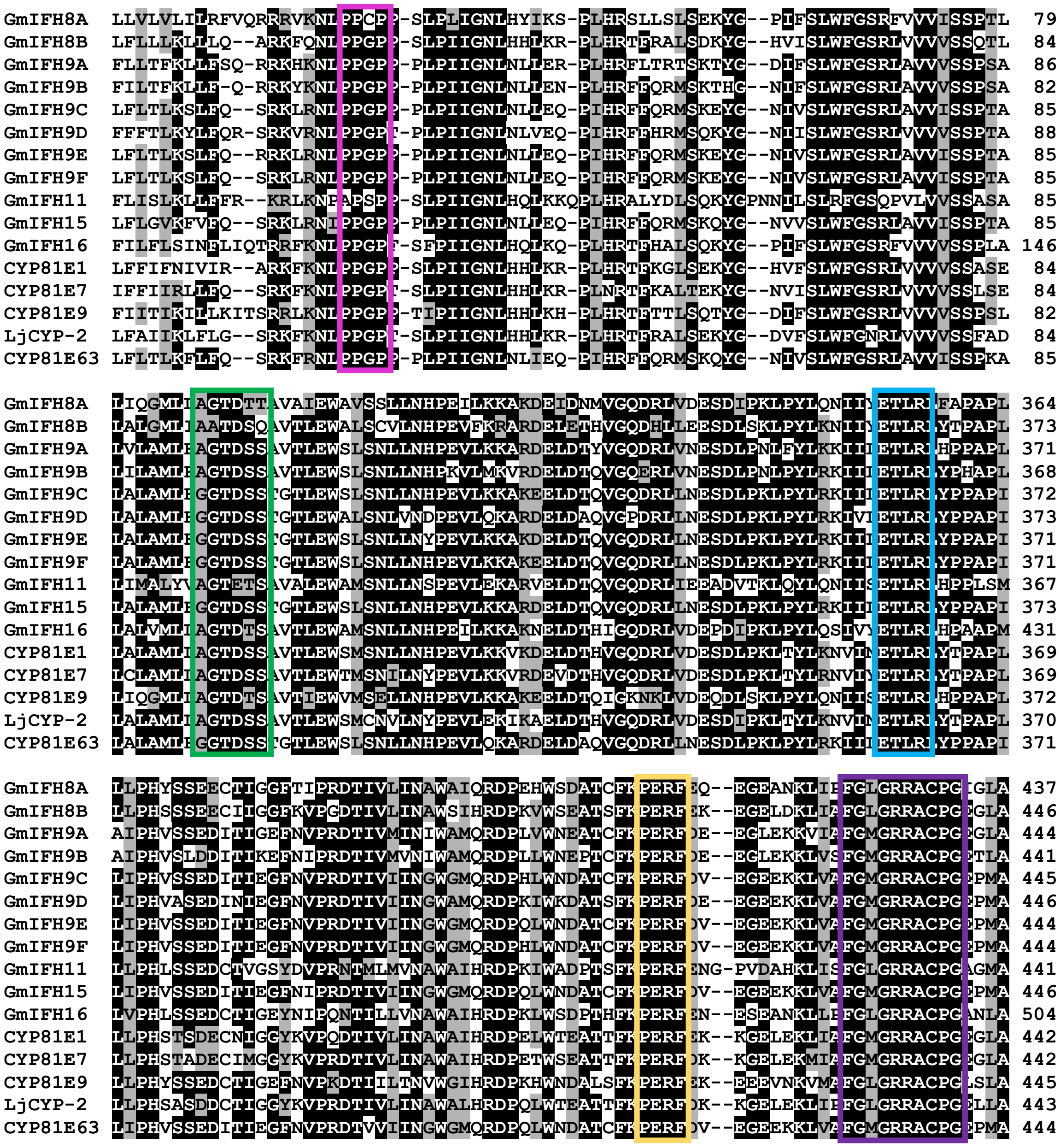


**Supplementary Figure 2.** Multiple sequence alignment of deduced amino acid sequences of candidate GmIFHs with characterized I2'H (CYP81E1, CYP81E7, CYP81E63, LjCYP-2) and I3'H (CYP81E9) from other legumes. Identical and similar amino acid residues are indicated by black and gray shades, respectively. Five P450 motifs: the PPGP (pink), the oxygen-binding (green), the ETLR (blue), the PERF (yellow) and the heme-binding motifs (purple) are indicated by different colored rectangles. Only an abridged version of the alignment is shown.

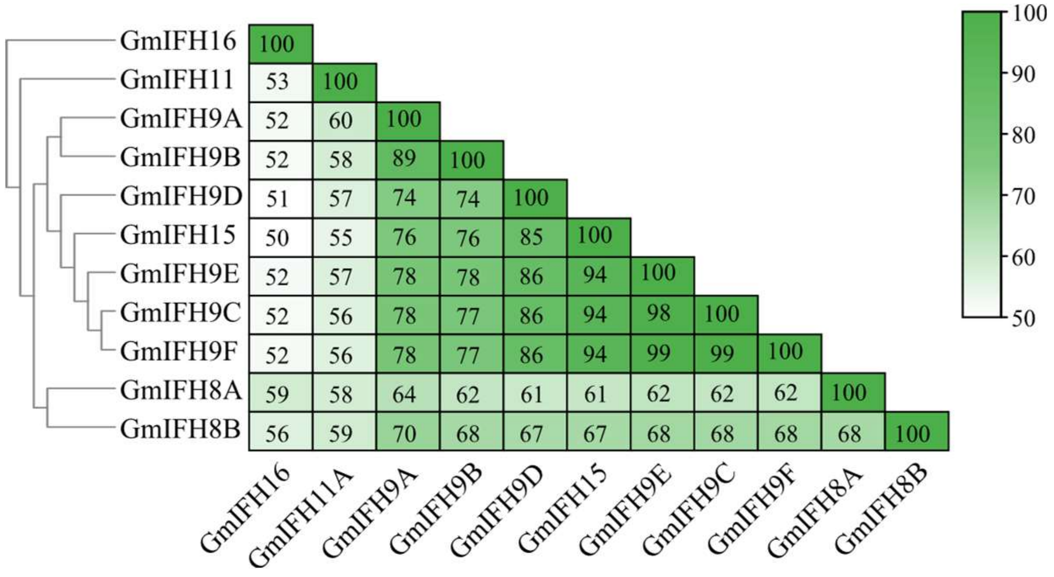


**Supplementary Figure 3.** GmIFH sequence identity. Sequence identity heatmap showing the pairwise percentage identity between GmIFH amino acid sequences. The X and Y axes indicate 11 identified GmIFHs. Identity scores are shown as a color-coded matrix, calculated by comparing every sequence to each other. Scale bar showing increase in sequence identity from white to green. Percent identity is also shown in each box.

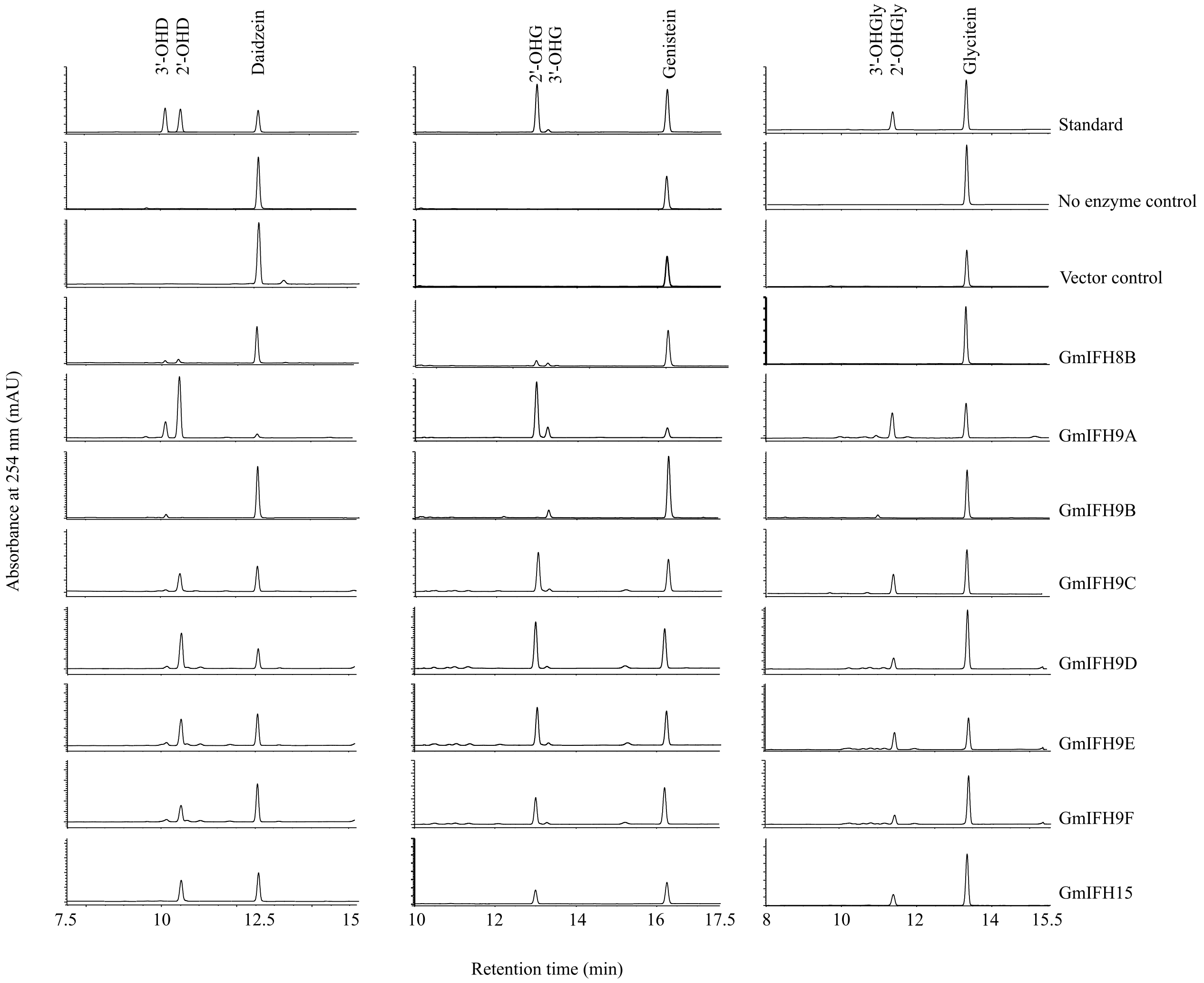


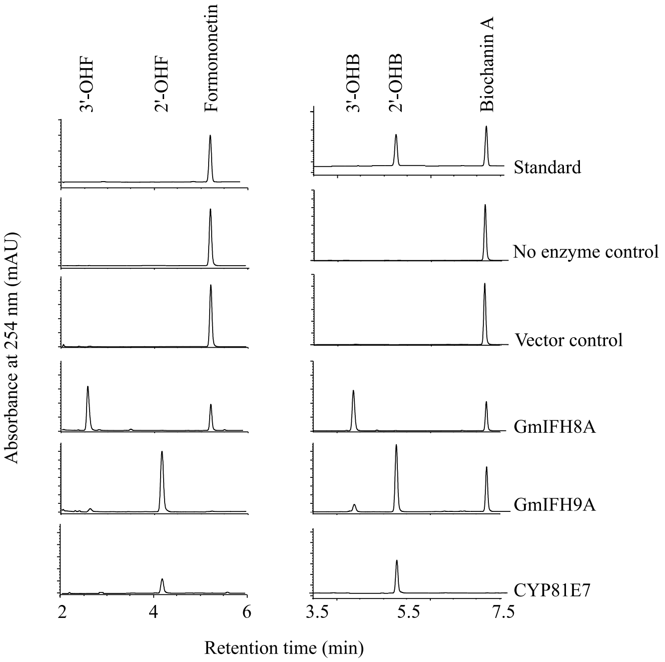


**Supplementary Figure 4**. Functional characterization of candidate GmIFHs in vitro. HPLC chromatogram showing the substrate and product peaks detected at 254 nm UV detector. Microsomal fraction containing candidate GmIFHs and LjCPR1 was incubated with different isoflavone aglycone substrates and NADPH followed by ethyl acetate extraction and analyzed by HPLC. Dotted lines indicate substrate and product peaks. 2'-OHD: 2'-hydroxydaidzein, 3'-OHD: 3'-hydroxydaidzein, 2'-OHG: 2'-hydroxygenistein, 3'-OHG: 3'-hydroxygenistein, 2'-OHGly: 2'-hydroxyglycitein, 3'-OHGly: 3'-hydroxyglycitein, F: Formononetin, 2'-OHF: 2'-hydroxyformononetin, 3'-OHF: 3'-hydroxyformononetin, B: Biochanin A, 2'-OHB: 2'-hydroxybiochanin A, 3'-OHB: 3'-hydroxybiochanin A. CYP81E7: *Medicago truncatula* I2'H.

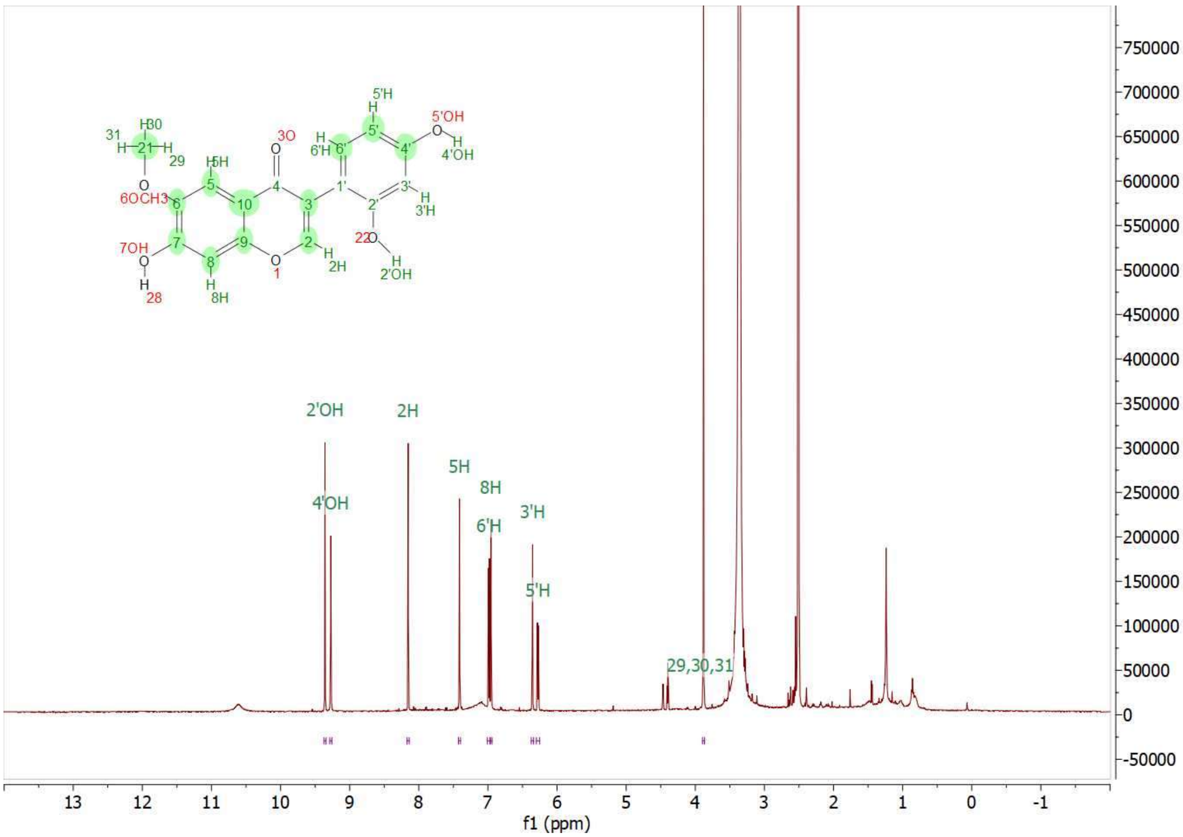


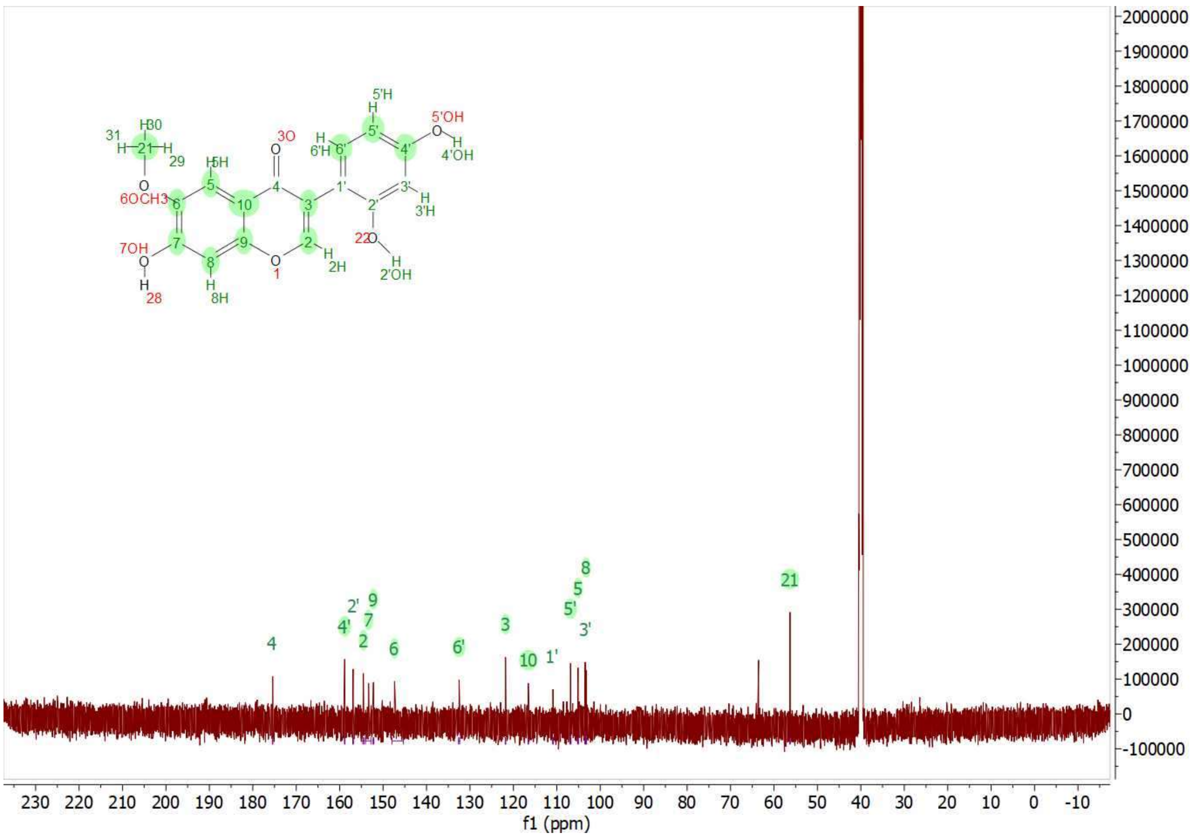


**Supplementary Figure 5:** 1H and 13C NMR spectra of 2'-hydroxyglycitein in DMSO-d6.

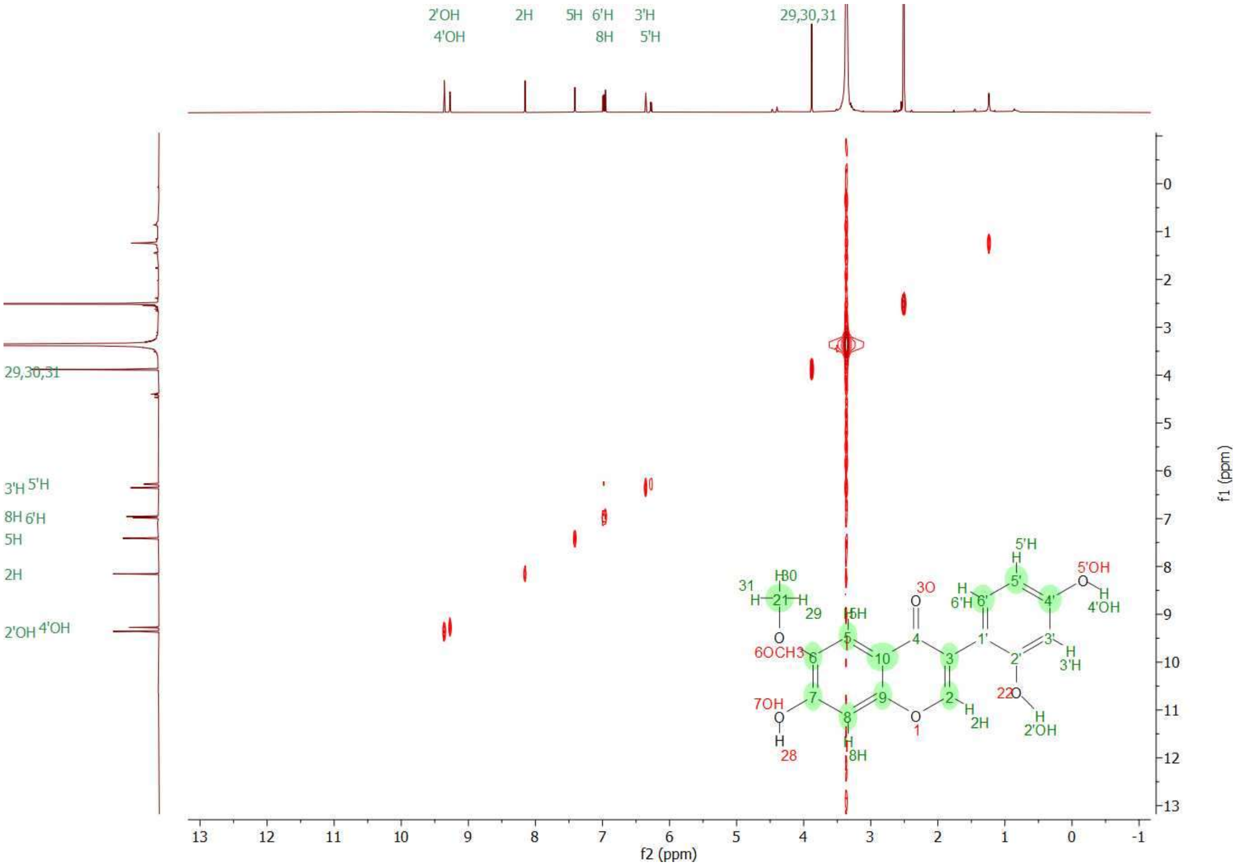


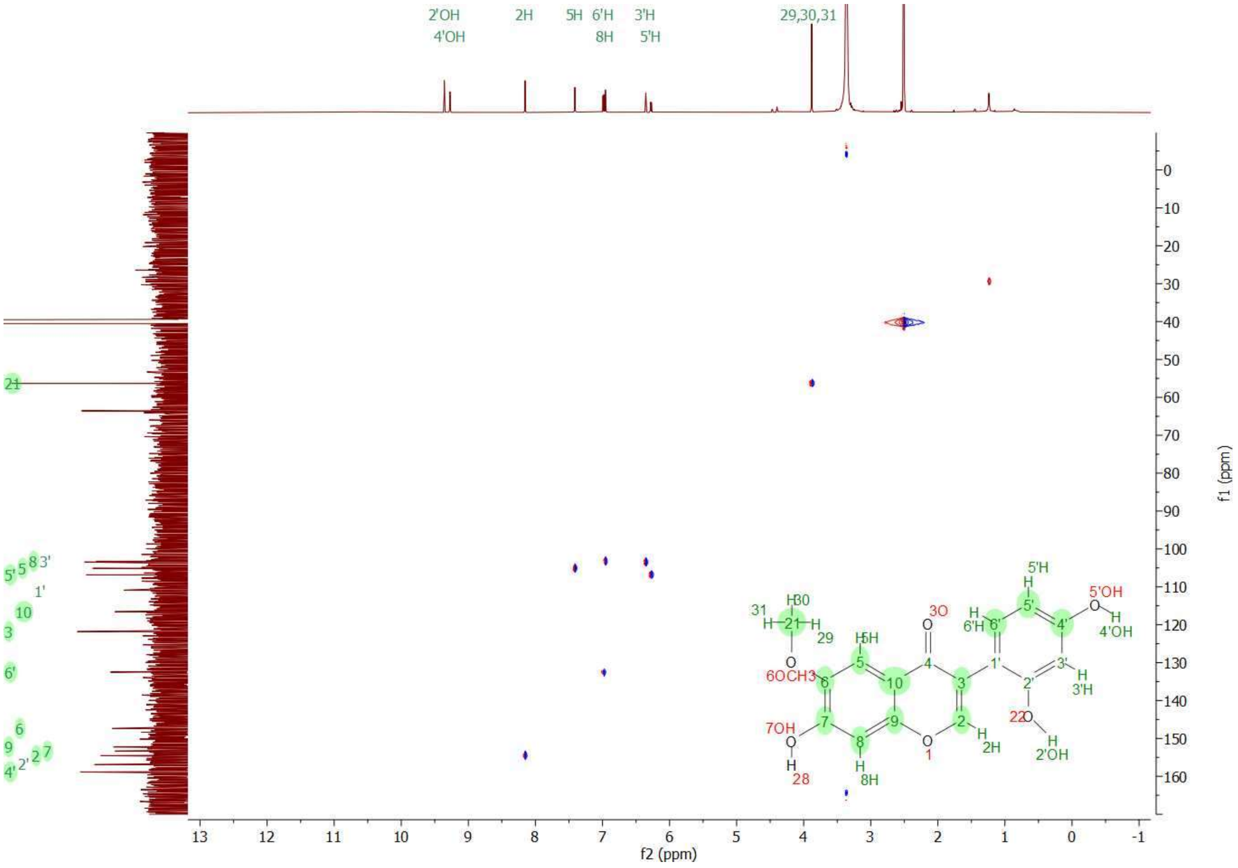


**Supplementary Figure 6:** COSY and HSQC NMR spectra of 2'-hydroxyglycitein in DMSO-d6.

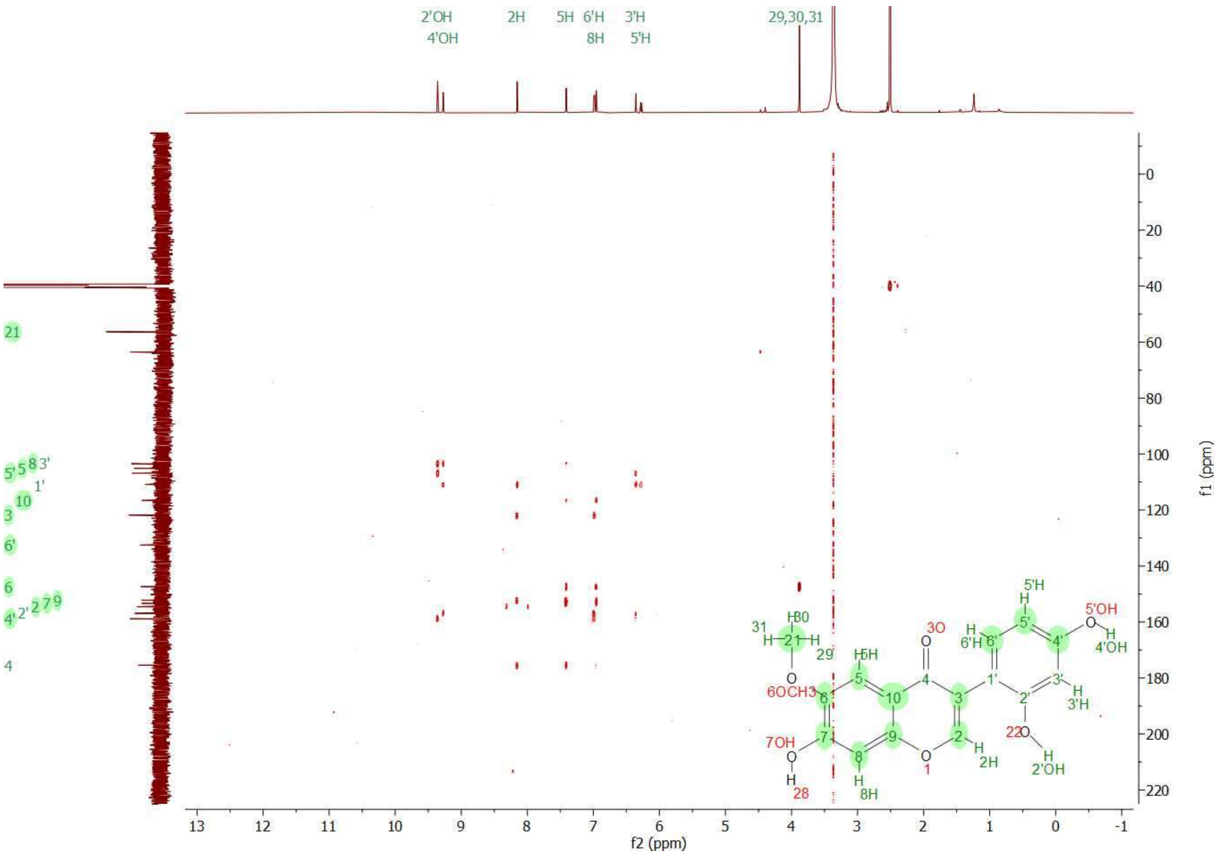


**Supplementary Figure 7:** HMBC NMR spectra of 2'-hydroxyglycitein in DMSO-d6**.**

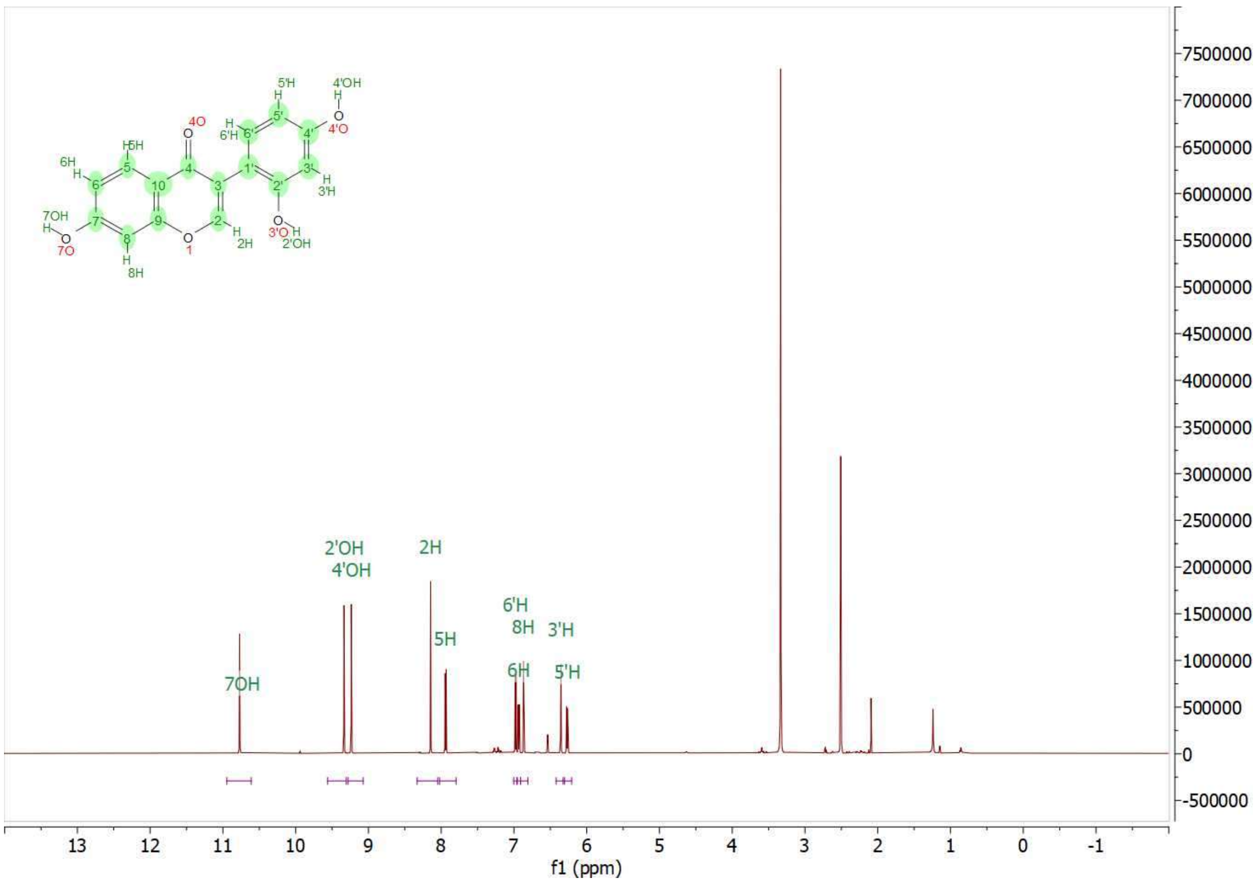


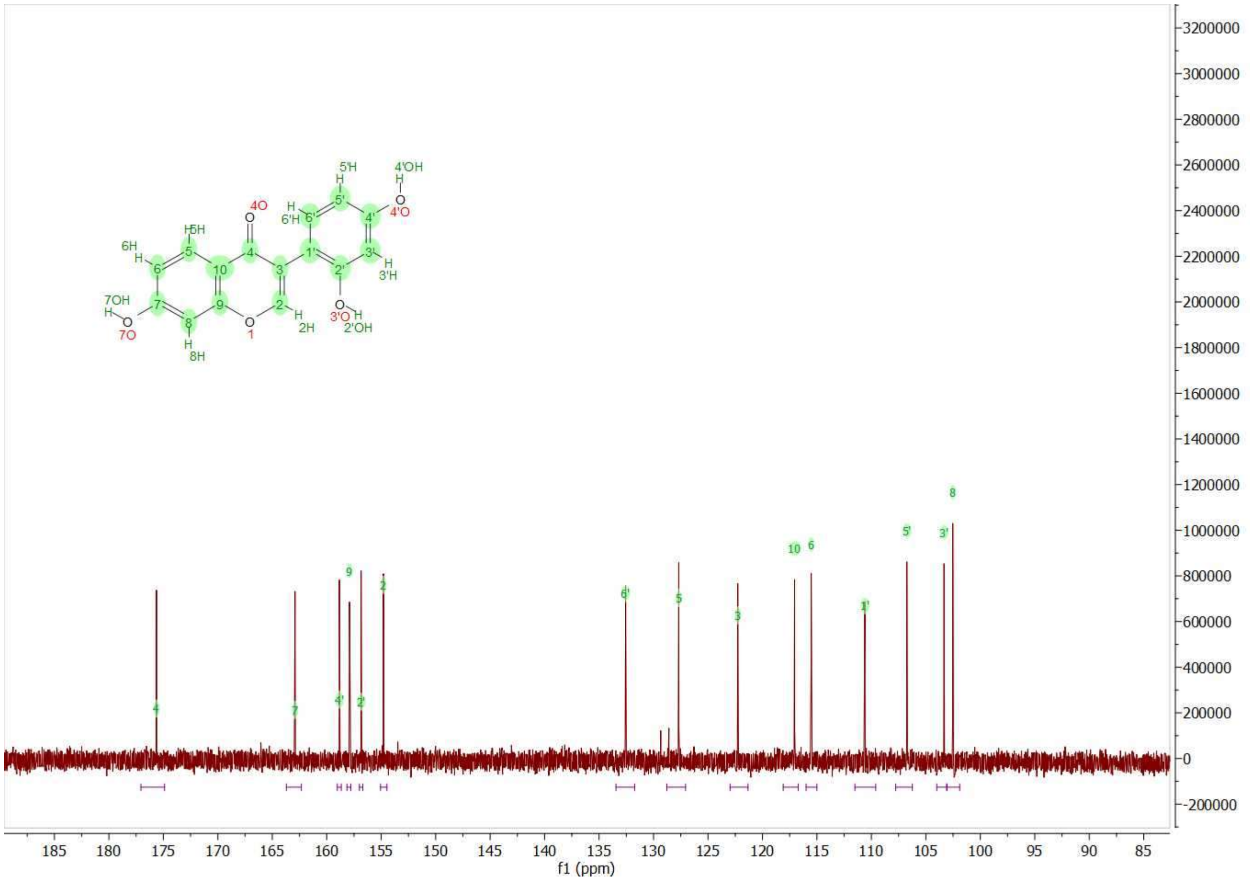


**Supplementary Figure 8:** 1H and 13C NMR spectra of 2'-hydroxydaidzein in DMSO-d6.

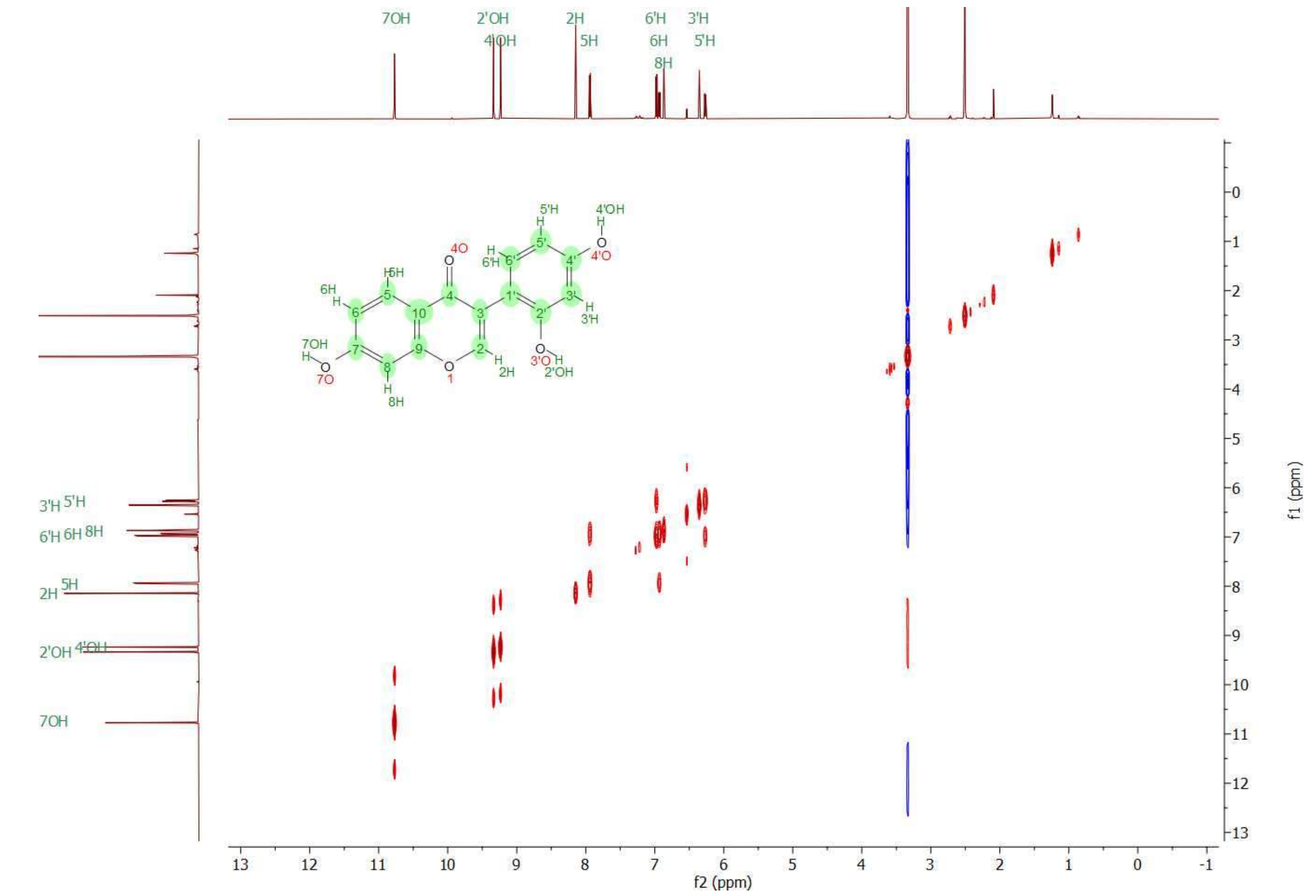


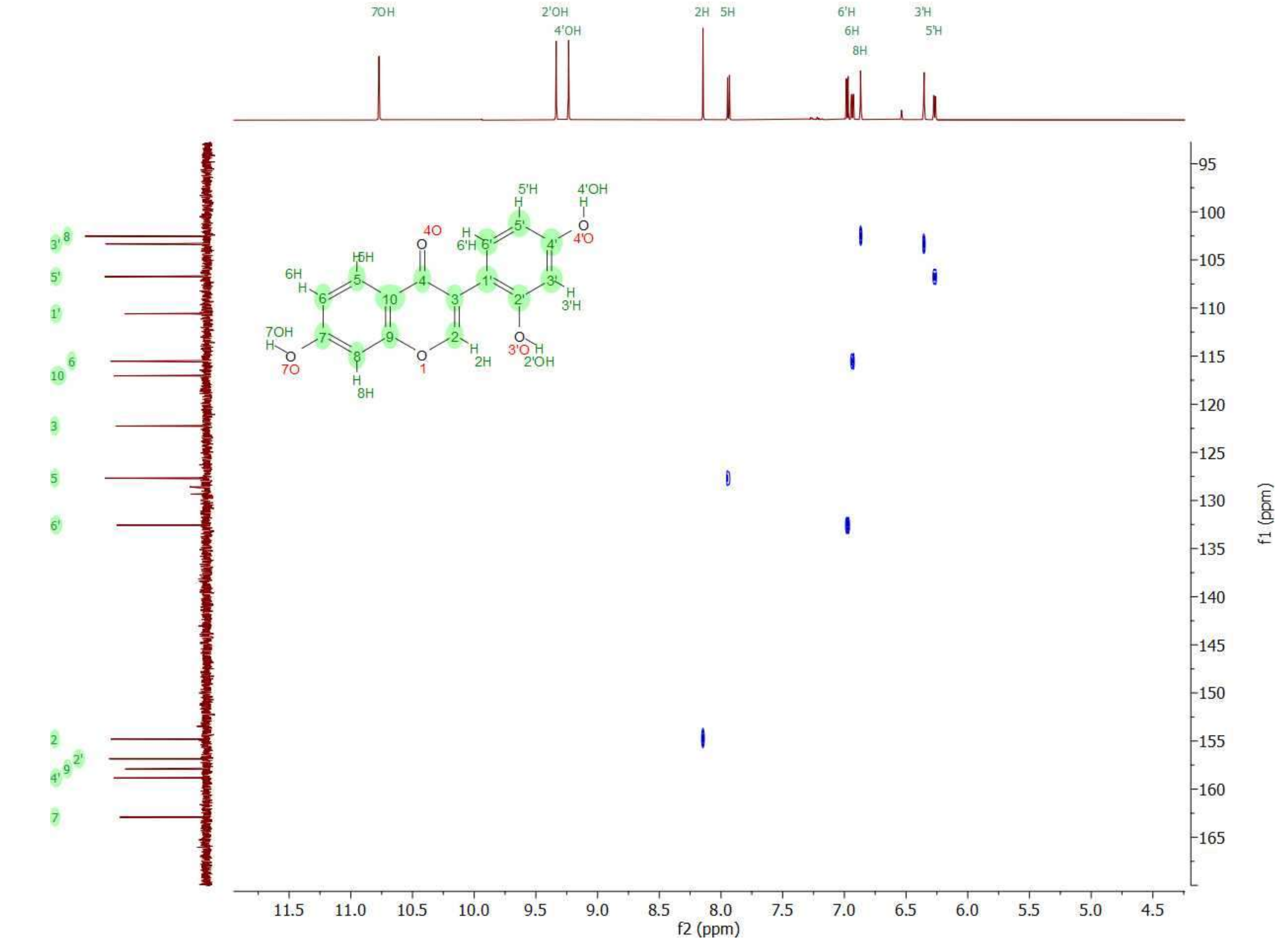


**Supplementary Figure 9:** COSY and HSQC NMR spectra of 2'-hydroxydaidzein in DMSO-d6.

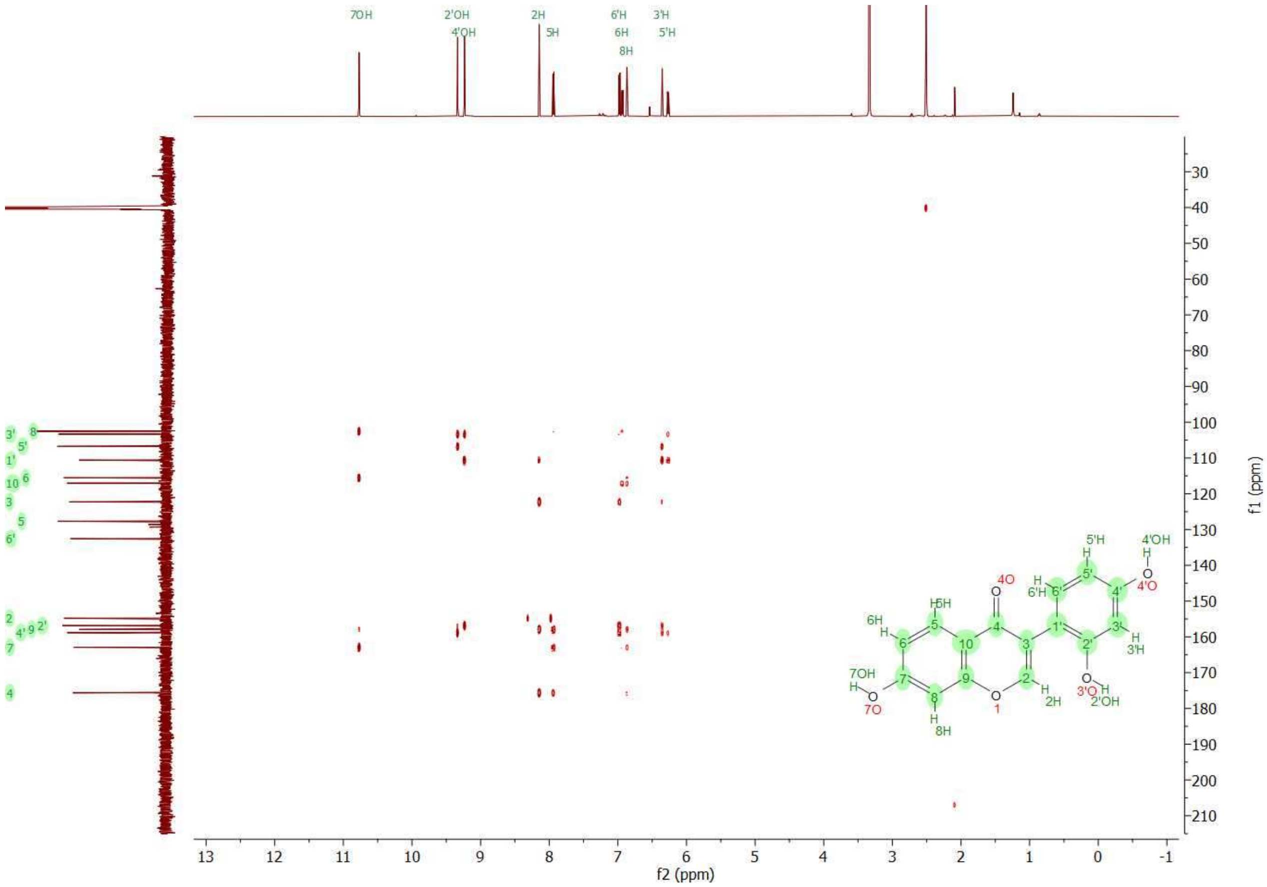


**Supplementary Figure 10:** HMBC NMR spectra of 2'-hydroxydaidzein in DMSO-d6.

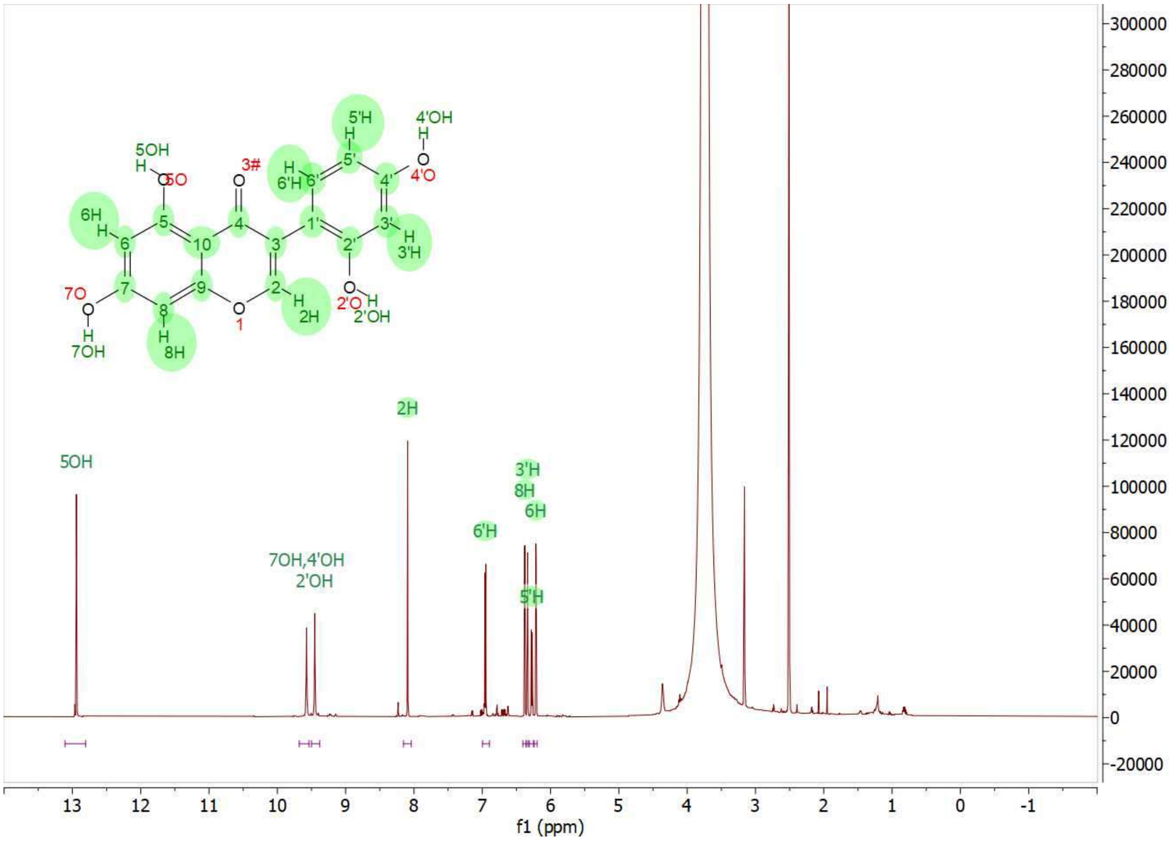


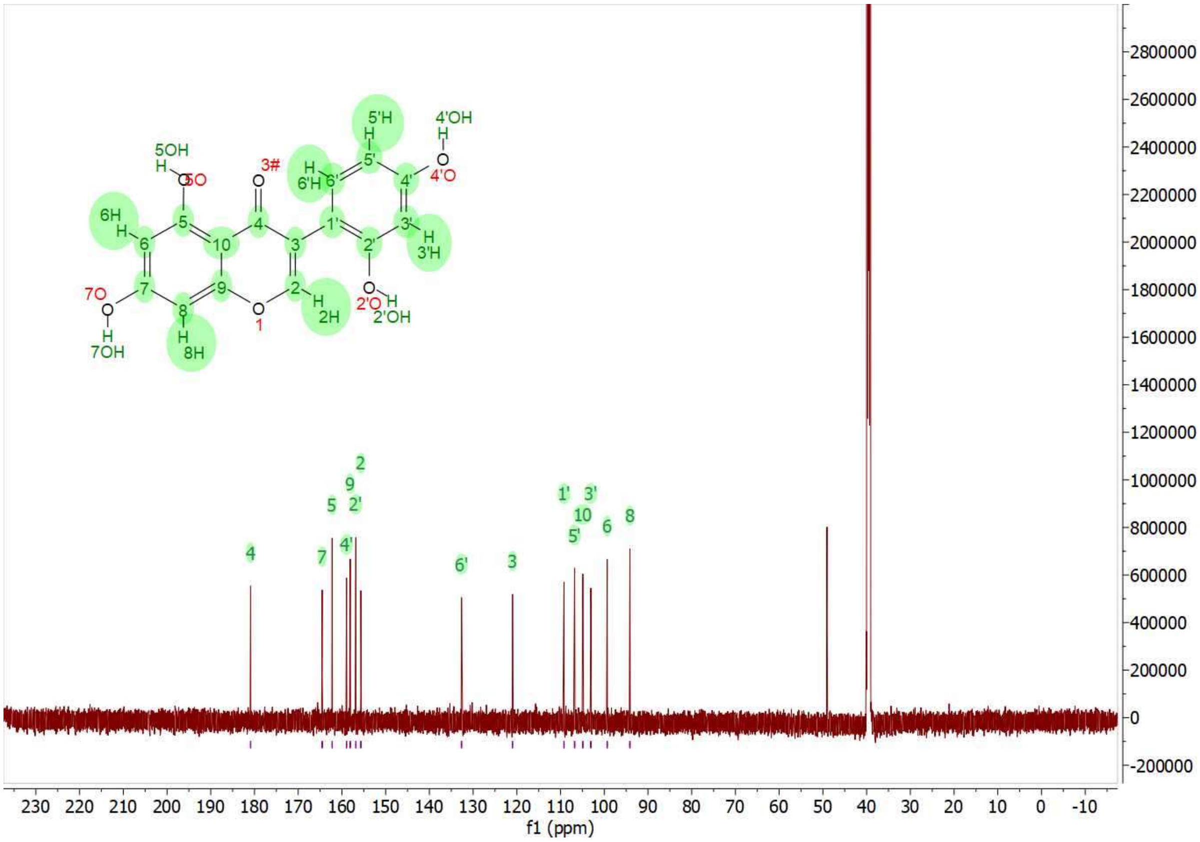


**Supplementary Figure 11**: 1H and 13C NMR spectra of 2'-hydroxygenistein in DMSO-d6.

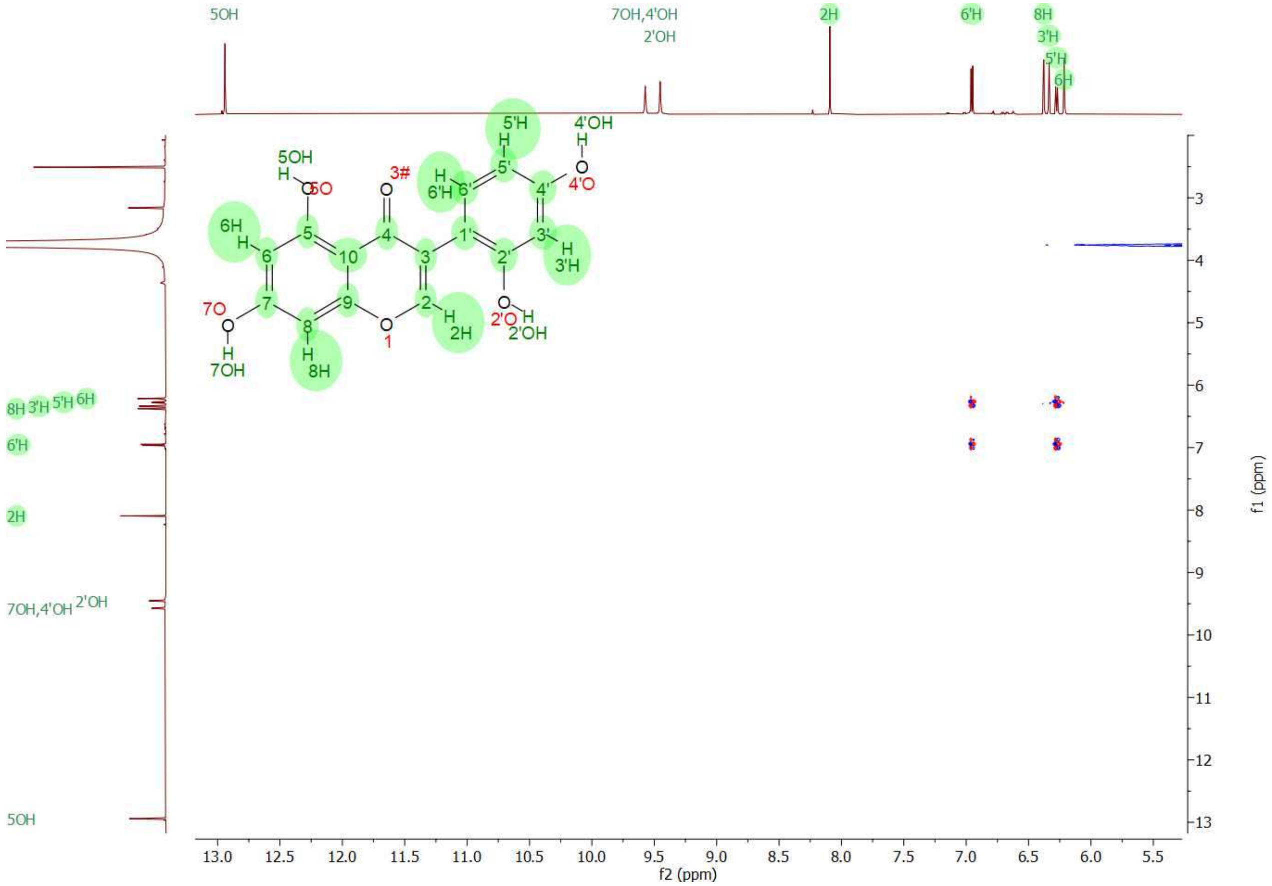


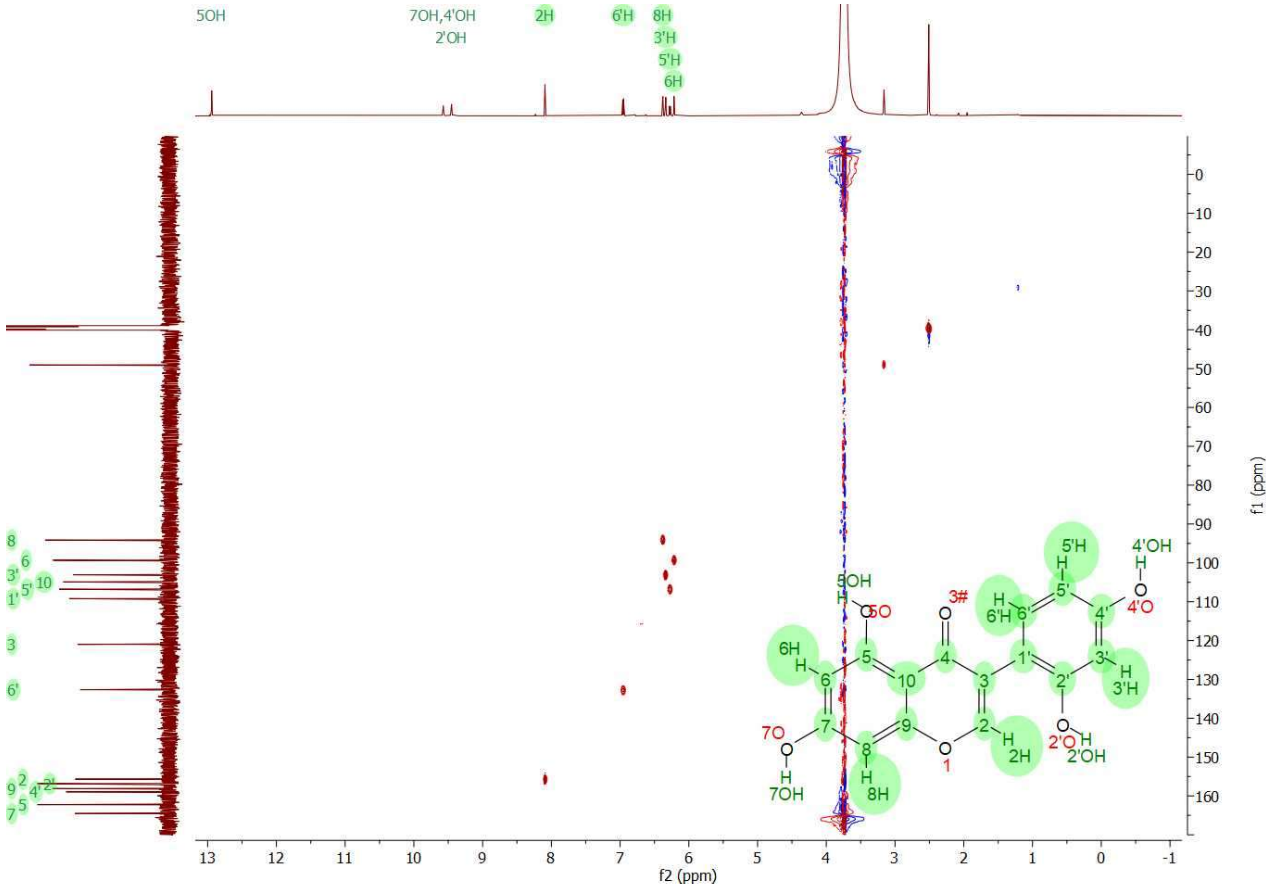


**Supplementary Figure 12:** COSY and HSQC NMR spectra of 2'-hydroxygenistein in DMSO-d6.

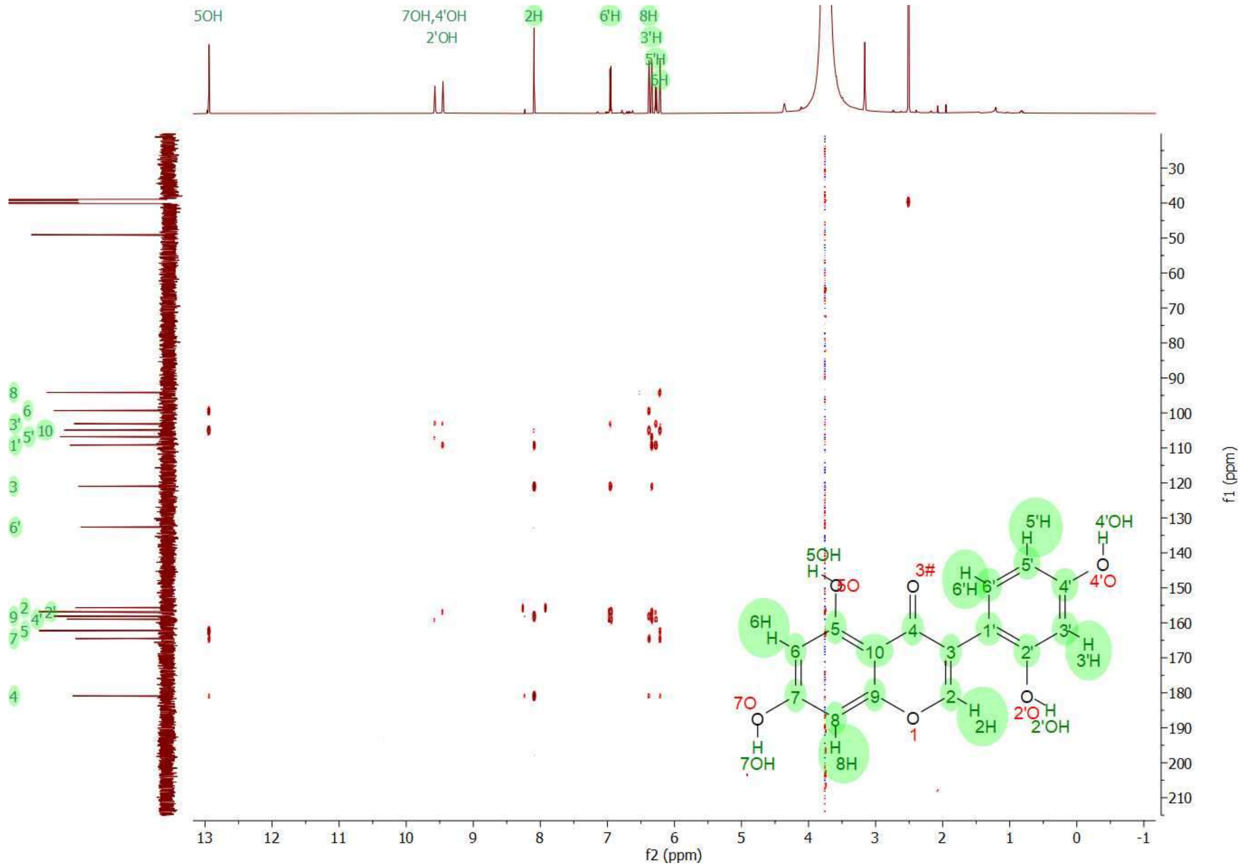


**Supplementary Figure 13**: HMBC NMR spectra of 2'-hydroxygenistein in DMSO-d6.

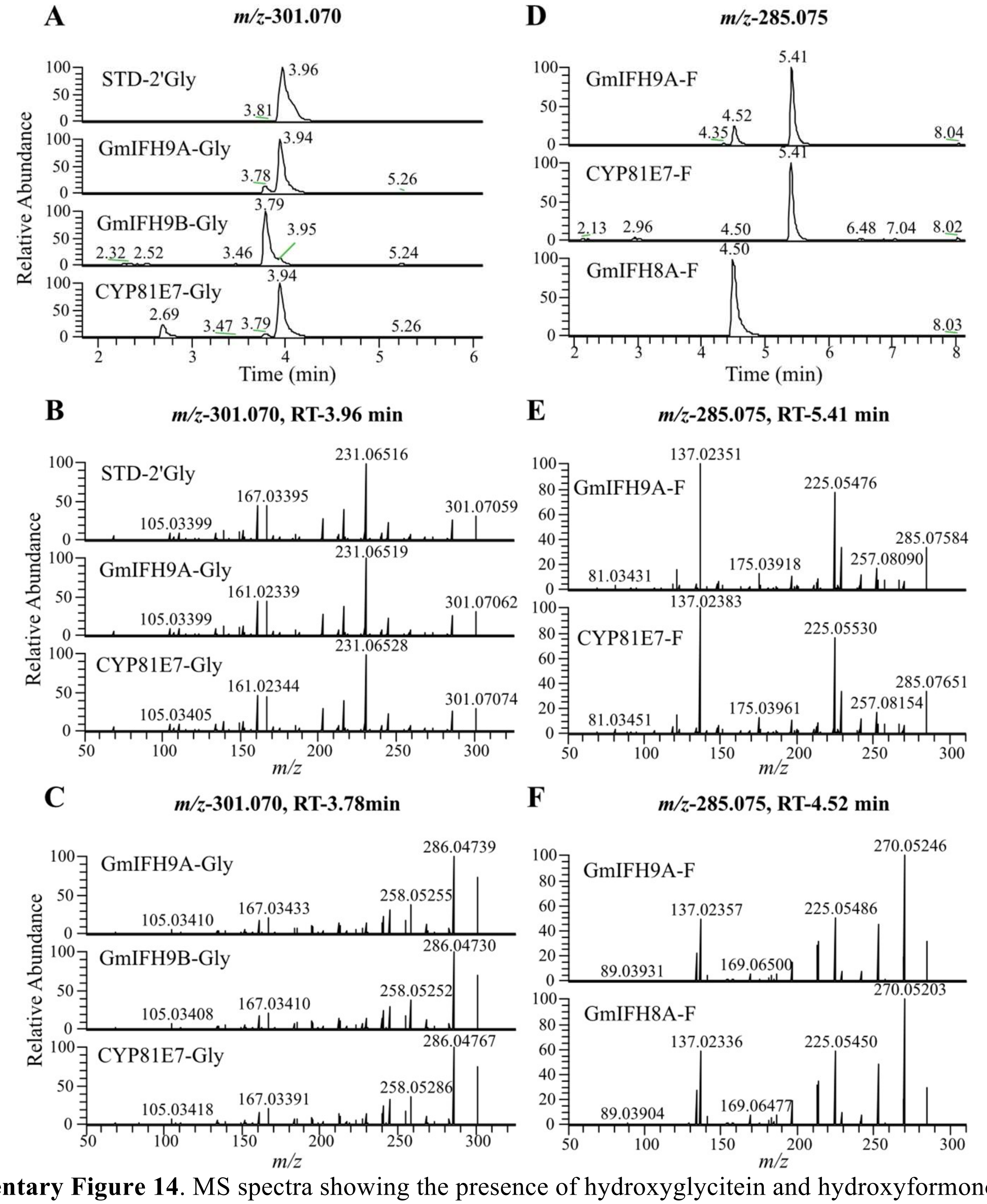


**Supplementary Figure 14**. MS spectra showing the presence of hydroxyglycitein and hydroxyformononetin in the GmIFH enzyme assay (A) Base peak at *m/z* 301.070 showing 2'-hydroxyglycitein (2'HGly) and 3'-hydroxyglycitein (3'HGly) at retention times (RT) of 3.96 and 3.78 min, respectively, in 2'HGly NMR verified compound standard (STD) and GmIFH9A, GmIFH9B and *Medicago truncatula* I2'H (CYP81E7) enzyme assay reactions with substrate glycitein (Gly). (B, C) MS/MS fragmentation of samples in panels A. (D) Base peak at *m/z* 285.075 showing 2'-hydroxyformononetin (2'HF) and 3'-hydroxyformononetin (3'HF) at retention times (RT) of 5.41 and 4.52 min, respectively, in GmIFH9A and GmIFH8A enzyme assay reactions incubated with substrate formononetin (F). *Medicago truncatula* I2'H (CYP81E7) enzyme assay reaction with substrate F used as a positive control for confirming 2'HF. (E, F) MS/MS fragmentation of samples in panel D.

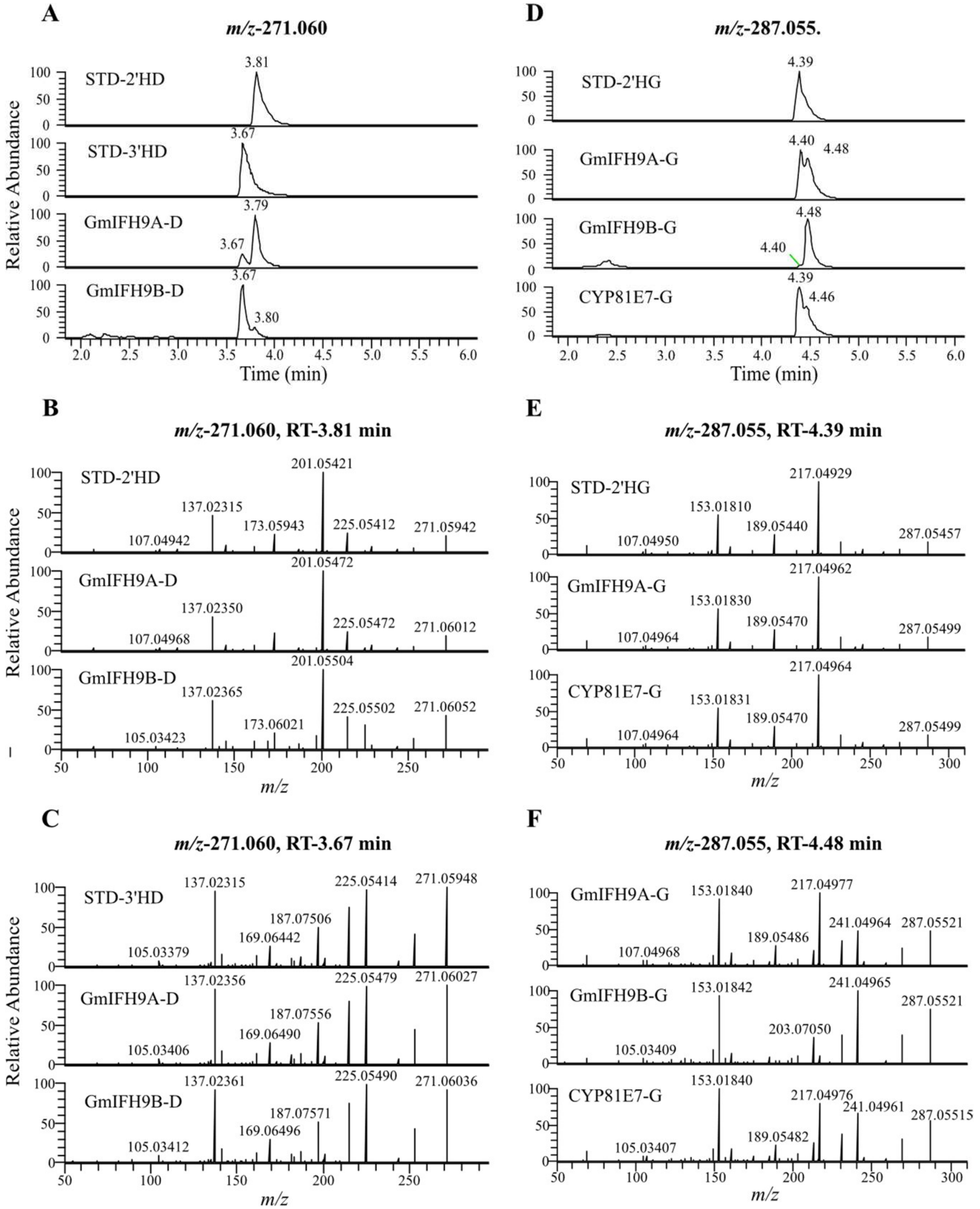


**Supplementary Figure 15.** MS spectra showing the presence of hydroxydaidzein and hydroxygenistein in the GmIFH enzyme assay. (A) Base peak at *m/z* 271.060 showing 2'-hydroxydaidzein (2'HD) and 3'-hydroxydaidzein (3'HD) at retention times (RT) of 3.81 and 3.67 min, respectively, in 2'HD, 3'HD standards and GmIFH9A and GmIFH9B enzyme assay reactions incubated with substrate daidzein (D), (B, C) MS/MS fragmentation of samples in panels A, (D) Base peak at *m/z* 287.055 showing 2'-hydroxygenistein (2'HG) and 3'-hydroxygenistein (3'HG) at retention times (RT) of 4.39 and 4.48 min, respectively, in 2'HG NMR verified compound standard (STD) and GmIFH9A and GmIFH9B enzyme assay reactions incubated with substrate genistein (G) The *Medicago truncatula* I2′H/CYP81E7 enzyme assay with genistein was used as a positive control for 2′-hydroxygenistein identification, (E, F) MS/MS fragmentation of samples in panel D.

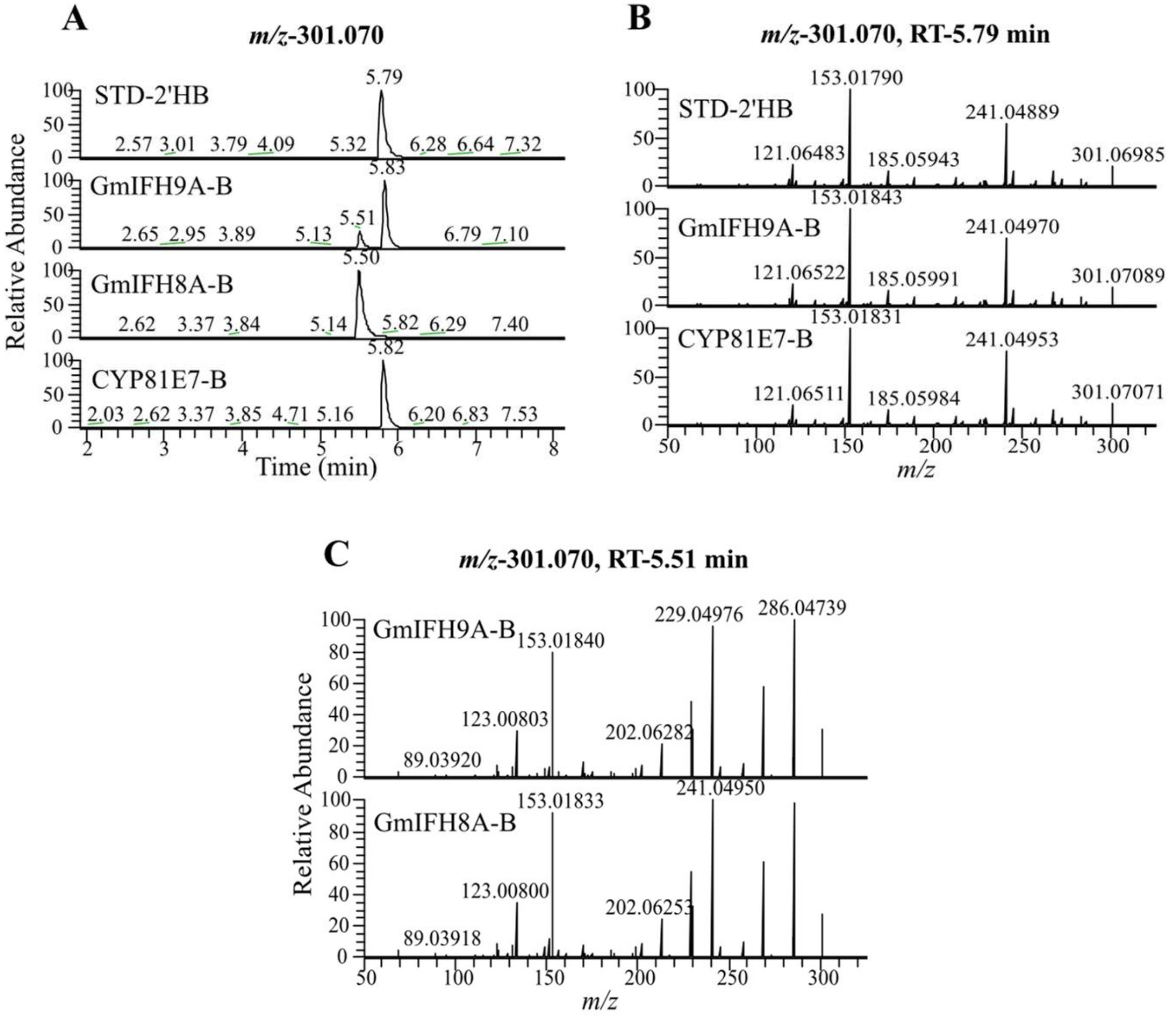


**Supplementary Figure 16**. MS spectra showing the presence of hydroxybiochanin A in the GmIFH enzyme assay. (A) Base peak at *m/z* 301.070 showing 2'-hydroxybiochanin A (2'HB) and 3'-hydroxybiochanin A (3'HB) at retention times (RT) of 5.79 and 5.51 min, respectively, in 2'HB standard and GmIFH9A and GmIFH8A enzyme assay reactions with substrate biochanin A (B). *Medicago truncatula* I2'H (CYP81E7) enzyme assay reaction with substrate B used as a positive control for confirming 2'HB. (B, C) MS/MS fragmentation of samples in panels A.

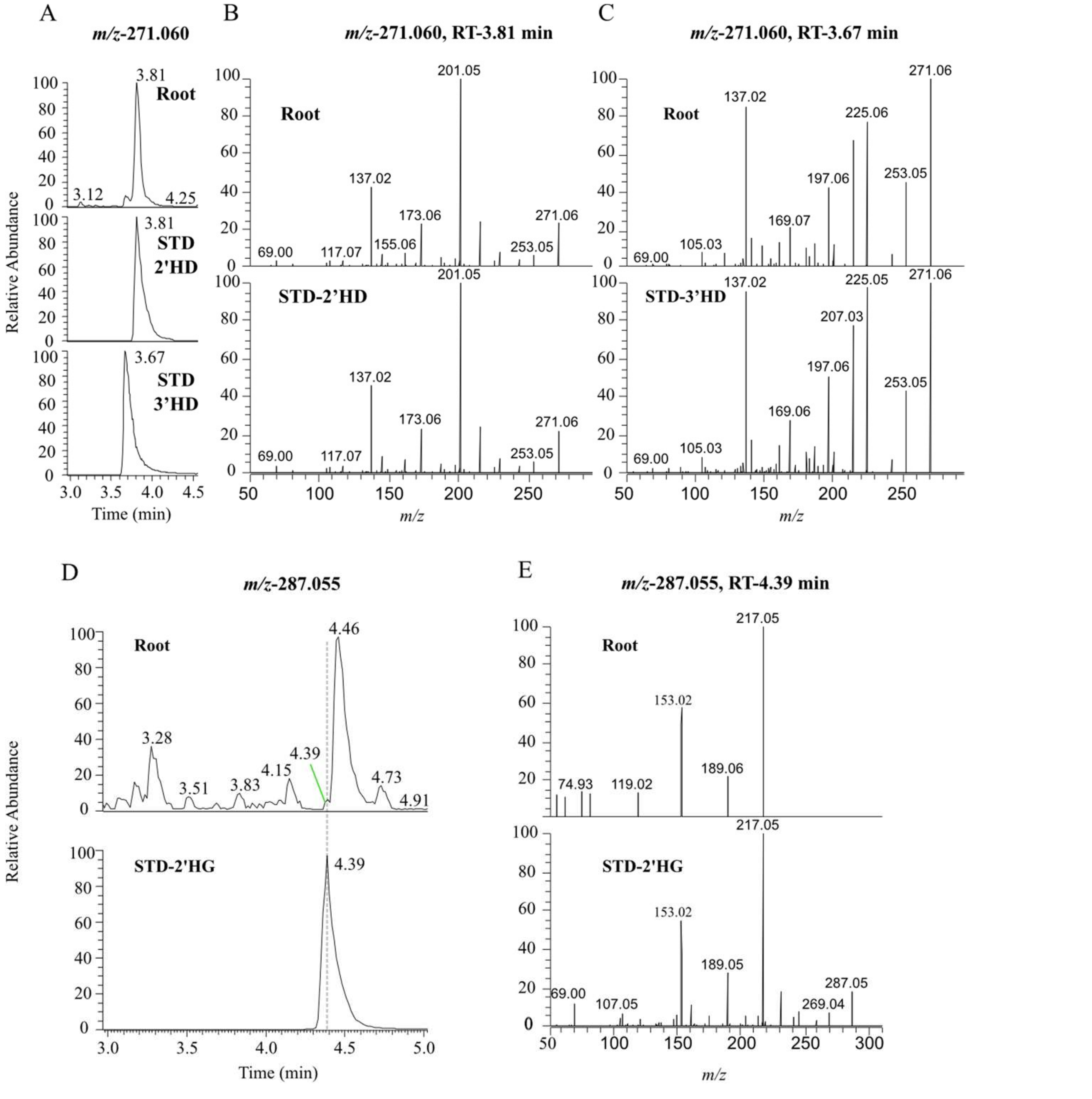


**Supplementary Figure 17.** Confirmation of hydroxydaidzein and hydroxygenistein in soybean roots. (A) Base peak at *m/z* 271.060 showing 2'-hydroxydaidzein (2'HD) and 3'-hydroxydaidzein (3'HD) at retention times (RT) of 3.71 and 3.57 min, respectively, in soybean root extract and authentic standards of 2'HD and 3'HD. (B, C) MS/MS fragmentation of samples in panels A, (D) Base peak at *m/z* 287.055 showing 2'-hydroxygenistein (2'HG) at RT 4.30 min in soybean root extract and the 2'HG NMR verified compound standard, (E) MS/MS fragmentation of the sample in panel D.
