## Supplementary Table 2 for "Multi-Substrate Specificity of Isoflavone hydroxylases (GmIFH) Drive Isoflavonoid Diversification in Soybean"

Supplementary Table 2. Characterized CYP81 proteins in different plant species.

| **CYP** | **Organism** | **Pathway involved** | **Reference** |
| --- | --- | --- | --- |
| CYP81E9 | *Medicago truncatula* | Medicarpin biosynthesis (Isoflavonoid) | (Liu et al., 2003) |
| CYP81E7 | *Medicago truncatula* | Medicarpin biosynthesis (Isoflavonoid) | (Liu *et al.*, 2003) |
| CYP81E1 | *Glycyrrhiza echinata* | Phytoalexin biosynthesis (Isoflavonoid) | (Akashi et al., 1998) |
| CYP81E63 | *Pueraria mirifica* | Miroestrol biosynthesis (Isoflavonoid) | (Suntichaikamolkul et al., 2022) |
| LjCYP-2 | *Lotus japonicus* | Vestitol biosynthesis (Isoflavonoid) | (Shimada et al., 2000) |
| CYP81Q1 | *Sesamum indicum* | Sesamin biosynthesis (lignans) | (Koyama et al., 2022) |
| CYP81Q3 | *Sesamum alatum* | Sesamin biosynthesis (lignans) | (Ono et al., 2018) |
| CYP81AM1 | *Tripterygium wilfordii* | Abietane diterpenoids biosynthesis | (Wang et al., 2021) |
| CYP81Q59 | *Cucumis melo* | Cucurbitacins biosynthesis (tetracyclic triterpenoids) | (Zhou et al., 2016) |
| CYP81Q58 | *Cucumis sativus* | Cucurbitacins biosynthesis (tetracyclic triterpenoids) | (Zhou *et al.*, 2016) |
| CYP81AQ19 | *Momordica charantia* | Triterpenoid biosynthesis | (Takase et al., 2019) |
| SrCYP81 | *Stevia rebaudiana* | Steviol biosynthesis (diterpenes) | (Gold et al., 2018) |
| CYP81C16 | *Salvia miltiorrhiza* | Tanshinone biosynthesis (diterpenoids) | (Ren et al., 2023) |
| CYP81F1 | *Arabidopsis thaliana* | Indole Glucosinolate Modification | (Pfalz et al., 2011) |
| CYP81F2 | *Arabidopsis thaliana* | Indole Glucosinolate Modification | (Pfalz *et al.*, 2011) |
| CYP81F3 | *Arabidopsis thaliana* | Indole Glucosinolate Modification | (Pfalz *et al.*, 2011) |
| CYP81F4 | *Arabidopsis thaliana* | Indole Glucosinolate Modification | (Pfalz *et al.*, 2011) |
| CYP81B140 | *Plumbago zeylanica* | Plumbagin biosynthesis (naphthoquinone) | (Vasav et al., 2022) |
| CYP81B141 | *Plumbago zeylanica* | Plumbagin biosynthesis (naphthoquinone) | (Vasav *et al.*, 2022) |
| CYP81AN15 | *Erythroxylum coca* | Tropane alkaloid biosynthesis | (Chavez et al., 2022) |
| HcCYP81AA1 | *Hypericum calycinum* | Xanthone biosynthesis | (El-Awaad et al., 2016) |
| HpCYP81AA1 | *Hypericum perforatum* | Xanthone biosynthesis | (El-Awaad *et al.*, 2016) |
| HpCYP81AA2 | *Hypericum perforatum* | Xanthone biosynthesis | (El-Awaad *et al.*, 2016) |
| HpCYP81AA3 | *Hypericum perforatum* | Xanthone biosynthesis | (El-Awaad *et al.*, 2016) |
| CYP81B1s | *Helianthus tuberosus* | In-chain monohydroxylation of medium chain saturated fatty acids | (Cabello-Hurtado et al., 1998) |
| CYP81B1l | *Helianthus tuberosus* | In-chain monohydroxylation of medium chain saturated fatty acids | (Cabello-Hurtado *et al.*, 1998) |
| CYP81A37 | *Zea mays* | Zealexin pathway (sesquiterpenoid) | (Ding et al., 2020) |
| CYP81A38 | *Zea mays* | Zealexin pathway (sesquiterpenoid) | (Ding *et al.*, 2020) |
| CYP81A39 | *Zea mays* | Zealexin pathway (sesquiterpenoid) | (Ding *et al.*, 2020) |
| CYP81A6 | *Oryza sativa* | Herbicides metabolism | (Ha et al., 2022) |
| CYP81A68 | *Echinochloa crus-galli* | Herbicides metabolism | (Pan et al., 2022) |
| CYP81A12 | *Echinochloa phyllopogon* | Herbicides metabolism | (Ha *et al.*, 2022) |
| CYP81A21 | *Echinochloa phyllopogon* | Herbicides metabolism | (Ha *et al.*, 2022) |
| CYP81A9 | *Zea mays* | Herbicides metabolism | (Brazier Hicks et al., 2022) |
| CYP81A2 | *Zea mays* | Herbicides metabolism | (Brazier Hicks *et al.*, 2022) |
| CYP81A16 | *Zea mays* | Herbicides metabolism | (Brazier Hicks *et al.*, 2022) |
| CYP81A15 | *Echinochloa phyllopogon* | Herbicides metabolism | (Dimaano et al., 2020) |
| CYP81A18 | *Echinochloa phyllopogon* | Herbicides metabolism | (Dimaano *et al.*, 2020) |
| CYP81A24 | *Echinochloa phyllopogon* | Herbicides metabolism | (Dimaano *et al.*, 2020) |
| CYP81A14 | *Echinochloa phyllopogon* | Herbicides metabolism | (Dimaano *et al.*, 2020) |
