## Supplementary Table 3 for "Multi-Substrate Specificity of Isoflavone hydroxylases (GmIFH) Drive Isoflavonoid Diversification in Soybean"

Supplementary Table 3. List of Primers used for gene cloning

| **Gene name** | **Primer name** | **Sequence (5' to 3')** |
| --- | --- | --- |
| GmIFH8A | GmIFH8AGWF | GGGGACAAGTTTGTACAAAAAAGCAGGCTTCATGGGTCCTTTGTTGTACTC |
|  | GmIFH8AGWR | GGGGACCACTTTGTACAAGAAAGCTGGGTATTAGGTACCAAATATAGCTAATT |
| GmIFH8B | GmIFH8BGWF | GGGGACAAGTTTGTACAAAAAAGCAGGCTTCATGGAGATCCTTTCCTTCTTG |
|  | GmIFH8BGWR | GGGGACCACTTTGTACAAGAAAGCTGGGTATTATTTCCCAAGCCTGTTGATG |
| GmIFH9A | GmIFH9AGWF | GGGGACAAGTTTGTACAAAAAAGCAGGCTTCATGGAAATGTTTCCCTTGTTGC |
|  | GmIFH9AGWR | GGGGACCACTTTGTACAAGAAAGCTGGGTATTACTTCAGATTAATCTTGTTC |
| GmIFH9B | GmIFH9BGWF | GGGGACAAGTTTGTACAAAAAAGCAGGCTTCATGTTTCCCTTGTTGCCTTG |
|  | GmIFH9BGWR | GGGGACCACTTTGTACAAGAAAGCTGGGTATTATTTCAAATTAATCTTG |
| GmIFH9C | GmIFH9CGWF | GGGGACAAGTTTGTACAAAAAAGCAGGCTTCATGGGAATGTTGTTGTTGTTGTC |
|  | GmIFH9CGWR | GGGGACCACTTTGTACAAGAAAGCTGGGTATTAAACTCCAATTTTAGTGGCG |
| GmIFH9D | GmIFH9DGWF | GGGGACAAGTTTGTACAAAAAAGCAGGCTTCATGACAGTAATAACAATGCCTCC |
|  | GmIFH9DGWR | GGGGACCACTTTGTACAAGAAAGCTGGGTATTAATAACTTCCAACTTTGC |
| GmIFH9E | GmIFH9EGWF | GGGGACAAGTTTGTACAAAAAAGCAGGCTTCATGGGAATGTTGTTGTTGTTGTT |
|  | GmIFH9EGWR | GGGGACCACTTTGTACAAGAAAGCTGGGTATCAAATTTTAGTGGCAAGTGGGC |
| GmIFH9F | GmIFH9FGWF | GGGGACAAGTTTGTACAAAAAAGCAGGCTTCATGGGAATGTTGTTGTTGTTGTT |
|  | GmIFH9FGWR | GGGGACCACTTTGTACAAGAAAGCTGGGTATTAAACTCCAATTTTAGTGG |
| GmIFH11 | GmIFH11GWF | GGGGACAAGTTTGTACAAAAAAGCAGGCTTCATGGAAGGCAACCTTATC |
|  | GmIFH11GWR | GGGGACCACTTTGTACAAGAAAGCTGGGTATTAGAAAATCTTGCTAATGATC |
| GmIFH15 | GmIFH15GWF | GGGGACAAGTTTGTACAAAAAAGCAGGCTTCATGGGAATGTTGTTGGTGGTG |
|  | GmIFH15GWR | GGGGACCACTTTGTACAAGAAAGCTGGGTATTAAATTCCAATTTTAGTGG |
| GmIFH16 | GmIFH16GWF | GGGGACAAGTTTGTACAAAAAAGCAGGCTTCATGACTCCTTTTTACTTCC |
|  | GmIFH16GWR | GGGGACCACTTTGTACAAGAAAGCTGGGTACTAGAAGTTATAAATATCCTTAA |
