## Supplementary Table 4 for "Multi-Substrate Specificity of Isoflavone hydroxylases (GmIFH) Drive Isoflavonoid Diversification in Soybean"

**Supplementary Table 4.** Peptides detected in microsomal fraction of yeast expressing recombinant GmIFHs.

| **Sample** | **Peptide sequence (N- to C- terminal)** |
| --- | --- |
| GmIFH8A | LVDESDIPKLPYLQNIIYETLR |
| GmIFH8B | LFDFENLEKR |
| GmIFH9A | IILETLR |
| GmIFH9B | IILETLR |
| GmIFH9C | IILETLR |
| GmIFH9D | IVLETLR |
| GmIFH9E | IILETLR |
| GmIFH9F | IILETLR |
| GmIFH11 | LQYLQNIISETLR |
| GmIFH15 | IILETLR |
| GmIFH16 | RTDAFLQGLIDQHR |
